# Dissociable control of motivation and reinforcement by distinct ventral striatal dopamine receptors

**DOI:** 10.1101/2023.06.27.546539

**Authors:** Juan Enriquez-Traba, Hector E Yarur-Castillo, Rodolfo J Flores, Tenley Weil, Snehashis Roy, Ted B Usdin, Christina T LaGamma, Miguel Arenivar, Huikun Wang, Valerie S Tsai, Amy E Moritz, David R Sibley, Rosario Moratalla, Zachary Z Freyberg, Hugo A Tejeda

## Abstract

Dopamine release in striatal circuits, including the nucleus accumbens (NAc), tracks separable features of reward such as motivation and reinforcement. However, the cellular and circuit mechanisms by which dopamine receptors transform dopamine release into distinct constructs of reward remain unclear. Here, we show that dopamine D3 receptor (D3R) signaling in the NAc drives motivated behavior by regulating local NAc microcircuits. Furthermore, D3Rs co-express with dopamine D1 receptors (D1Rs), which regulate reinforcement, but not motivation. Paralleling dissociable roles in reward function, we report non-overlapping physiological actions of D3R and D1R signaling in NAc neurons. Our results establish a novel cellular framework wherein dopamine signaling within the same NAc cell type is physiologically compartmentalized via actions on distinct dopamine receptors. This structural and functional organization provides neurons in a limbic circuit with the unique ability to orchestrate dissociable aspects of reward-related behaviors that are relevant to the etiology of neuropsychiatric disorders.

## Main

Dopamine (DA) transmission is essential for reward function and its constituent features, including motivation and reinforcement. Motivation can be broadly defined as the internal process that activates and directs behavior, while reinforcement refers to the process in which the likelihood of a behavior is increased as a consequence of stimulus-response and action-outcome associations^1^. These separable constructs are coordinated to maximize reward outcomes^2–4^, and alterations in these reward sub-features are implicated in neuropsychiatric disorders, such as substance use and mood disorders^5–7^. A major integrative hub mediating motivation and reward-driven reinforcement is the nucleus accumbens (NAc), which receives DAergic projections from the ventral tegmental area (VTA)^8,9^. Disparate models have been proposed to explain the various components of reward that DA regulates in the NAc^10–14^. For instance, tonic and phasic patterns of DA neuron activity, and axonal control of DAergic terminals, contribute to dynamic fluctuations in DA concentrations that are hypothesized to ultimately underlie reinforcement, motivation, and vigor^12,15–22^. However, the specific mechanisms by which DA release is translated into cellular changes via dopamine receptors to drive dissociable features of reward function, such as motivation and reinforcement, remain unknown.

As in the dorsal striatum, medium spiny neurons (MSNs) of the NAc express high levels of DA receptors, including D1Rs and DA D2 receptors (D2Rs). Expression of D1Rs and D2Rs is segregated into non-overlapping MSN cell types, D1-MSNs and D2-MSNs, respectively^23,24^. DA-mediated activation of NAc D1Rs and D2Rs has been shown to produce dichotomous effects on D1- and D2-MSN function, respectively^25–29^. However, unlike its dorsal counterpart, the NAc is specifically enriched with the D3R^30–33^. Importantly, this specialized expression of D3Rs within ventral striatal circuits coincides with the specialized control of motivation and reinforcement by the NAc^8^. Moreover, D3R is a high-affinity DA receptor with a ten-fold greater DA affinity than D2Rs^34,35^, suggesting that it may serve as an additional conduit for the detection of tonic changes or dips in DA concentrations besides the D2R. Although D3R expression in the ventral striatum has been suggested, its role in regulating reward-related behaviors remains poorly understood. Important clues come from pharmacological manipulations of D3R signaling. Indeed, NAc D3R pharmacological antagonism has been shown to inhibit drug-seeking behavior^36,37^. However, an important limitation of these approaches is the relative inability of antagonists to precisely distinguish D3R from D2Rs. Furthermore, studies using either mice ectopically overexpressing D3R in all striatal MSNs or constitutive global knockout of D3Rs reported disrupted motivation and increased cocaine-seeking behavior, respectively^38,39^. However, striatum-wide D3R ectopic overexpression does not determine how endogenous NAc D3Rs contribute to reward function. Further, global knockout approaches do not distinguish contributions of NAc D3Rs versus D3Rs in other regions implicated in reward-seeking behavior^40,41^ or developmental compensation that may perturb reward function. Thus, limited tools to selectively manipulate the function of NAc D3Rs have hindered advances in our understanding of the specific contribution of D3Rs to distinct features underlying reward function. Interestingly, NAc D1-MSNs have been suggested to co-express D3R with D1R^41,42^, and this could provide NAc D1-MSNs with different modes by which DA may alter physiology and behavior. Because D1R and D3Rs possess different affinities for DA and engage opposite signaling effectors, these receptor-specific differences may translate into differential detection of DA concentrations and downstream effects on MSN function. We therefore hypothesized that D1-MSNs in the NAc may use distinct DA receptors for orchestrating dissociable reward-related functions.

Here we sought to address this hypothesis using a combination of anatomical tracing, slice electrophysiology and circuit-level manipulations of DA receptor function. In this study, we overcame technical limitations to selectively study NAc D3R fucntion by generating and characterizing a novel D3R conditional-knockout (cKO) mouse. Leveraging this line, we found that NAc D3R activity is necessary for motivated behavior, but not reinforcement, by acting presynaptically to inhibit NAc collateral transmission. Conversely, NAc D1Rs, with which D3Rs are highly co-expressed, promote reward- and aversion-driven reinforcement, but not motivation. Moreover, we demonstrated dissociable roles for D3R and D1R in regulating MSN synaptic physiology, which explained the dissociable roles in behavior. Our findings describe a novel framework by which DA signaling via D3R and D1R provides NAc D1-MSNs with the unique ability to regulate dissociable constructs underlying reward-related behaviors.

## Results

### Conditional knockout of NAc D3Rs results in motivational deficits

We first investigated whether NAc D3Rs regulate specific sub-features of reward-related behaviors. To overcome limitations associated with pharmacological antagonism and constitutive knockouts of D3Rs, we generated *Drd3* cKO mice (*Drd3*^fl/fl^) mice (Extended Data Fig. 1a) to selectively knockout D3R expression in the NAc. We achieved this aim by injecting Cre-expressing virus (AAV8-hSyn-GFP-Cre) into the NAc of *Drd3*^fl/fl^ mice (NAc-D3RcKO; Fig. 1a). Indeed, NAc-D3RcKO mice had decreased expression of *Drd3* mRNA as shown by fluorescent *in-situ* hybridization (Fig. 1b). Furthermore, we observed decreased *Drd3* mRNA expression as assessed by qPCR obtained from micro-dissected NAc. Importantly, no changes in *Drd1a* or *Drd2* expression were observed with NAc *Drd3* cKO, suggesting that this manipulation is specific to the D3R (Fig. 1c). We then assessed the impact of NAc D3R cKO in mediating reward function by first determining its role in motivation for exercise using wheel-running (Extended Data Fig. 1b), a rewarding stimulus for laboratory and feral rodents^43–45^. In this context, WT and *Drd3*^fl/fl^ control mice injected with Cre-expressing and GFP-expressing virus, respectively, displayed robust wheel running activity when exposed to a novel wheel during their inactive cycle (lights on), when animals should be at rest and locomotor activity is typically low (Fig. 1d). In contrast, NAc-D3RcKO mice had reduced wheel-running activity during this exposure to the novel running wheel in the inactive cycle (Fig. 1d), which dissipated prior to the onset of the active cycle (Extended Data Fig. 1c). However, running behavior triggered by the onset of the active cycle when lights turned off and did not differ between controls and NAc-D3RcKO mice (Extended Data Fig. 1c). Moreover, locomotor activity during the animals’ active cycle (lights off) in an open field was not impacted (Extended Data Fig. 1f). Intact locomotor activity during exploratory behavior in the open field and in response to diurnal shifts after the initial exposure to the running wheel, suggest that D3R cKO in the NAc is impacting the motivational value of running, not locomotor activity or running performance. Indeed, running behavior triggered by novel wheel exposure during the subjective daytime in rodents contains a strong motivational component^46–48^. Further, interference of running behavior is more sensitive during the acquisition of wheel running before habituation and experience^49^. With habituation to voluntary running, running behavior can be decoupled from enhancements in motivational state and driven by other factors, including but not limited to, habitual / stereotyped behavior, diurnal shifts, internal states, arousal-driven locomotor activity, and/or non-photic circadian entrainment^47,48^. In agreement with this notion, in rats habituated with running wheels, inactivation of the NAc does not impair running, unless motivation to run is increased by wheel deprivation^50^. Together, our results suggest that the initial motivation to engage in wheel running, but not locomotor activity or running performance, is decreased in NAc-D3RcKO mice.

**Figure 1.**
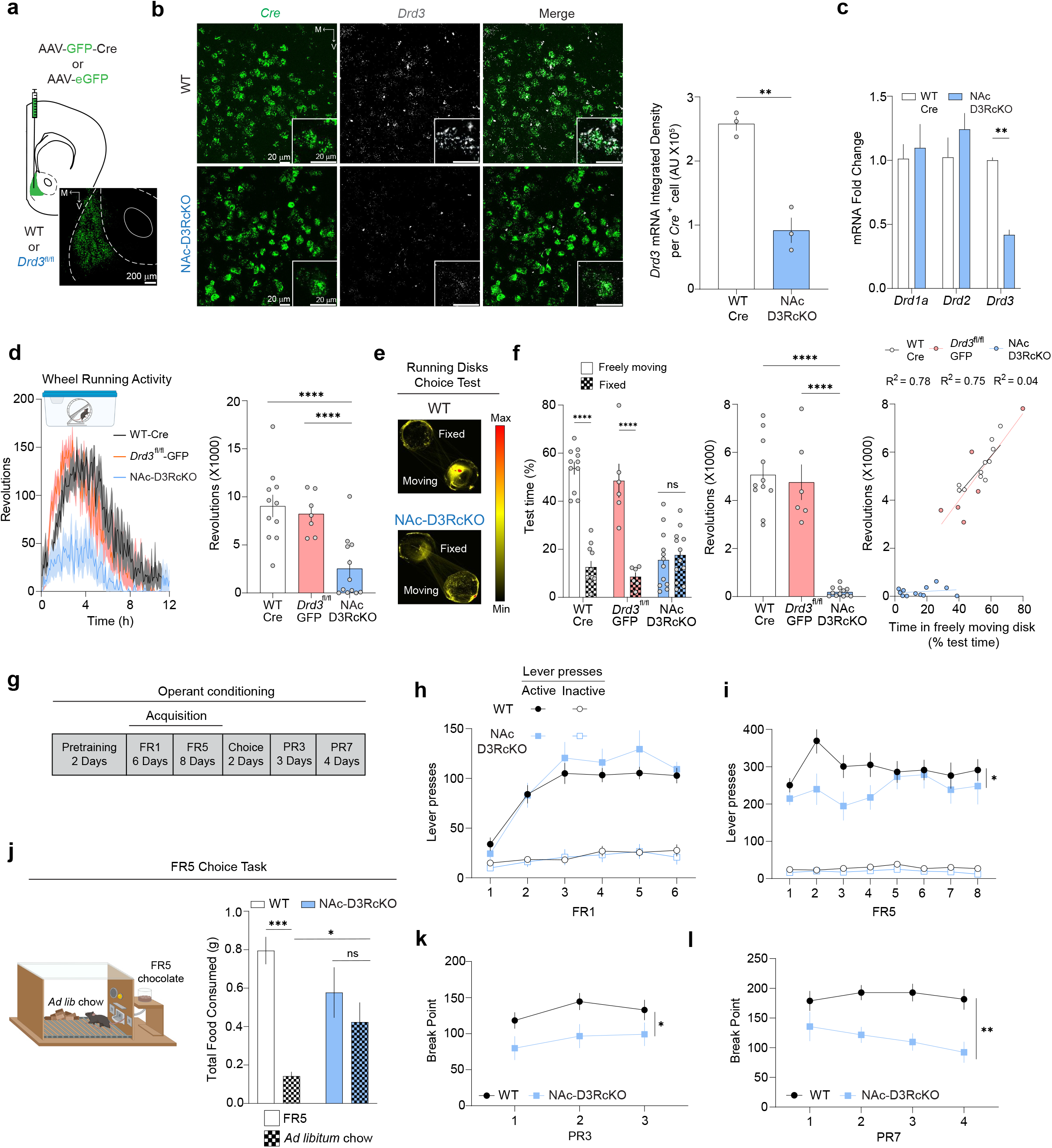
Conditional knockout of NAc D3Rs results in motivational deficits (see also Extended Data Figures Figure 1, 2 and 3) a. Experimental scheme (top) and representative image (bottom) of the NAc targeted with AAV-GFP-Cre or AAV-eGFP. b. (Left) Representative *in situ* hybridization images showing *Cre* (green) and *Drd*3 (white) mRNA expression in the NAc of WT (top) or NAc-D3RcKO mice (bottom). Insets depict higher-magnification images. (Right) Quantification of *Drd3* mRNA expression in *Cre*-positive neurons in the NAc of WT and NAc-D3RcKO mice. c. Quantitative real-time PCR analysis of *Drd1a*, *Drd2* and *Drd3* mRNA expression in the NAc of WT (white) and NAc-D3RcKO (blue) mice injected with Cre-expressing virus. d. (Left) Time-course of wheel-running activity in WT-Cre (black), *Drd3*^fl/fl^-GFP (red) and NAc-D3RcKO (blue) mice during the first 12 hrs of running wheel exposure. (Right) Quantification of revolutions across the 12-hr period. e. Representative occupancy heatmaps from WT (top) and NAc-D3RcKO (bottom) mice during running disks choice task. f. (Left) Quantification of time spent on the freely-moving and fixed disk. (Middle) Revolutions registered on the freely-moving disk for WT, *Drd3*^fl/fl^-GFP controls and NAc-D3RcKO mice. (Right) Spearman’s correlation between time spent in the freely-moving disk and revolutions. g. Timeline of operant conditioning experiment. h. Number of active and inactive lever presses of WT (black) and NAc-D3RcKO (blue) animals during FR1 acquisition sessions. i. Number of active and inactive lever presses of WT (black) and NAc-D3RcKO (blue) animals during FR5 (right) acquisition sessions. j. (Left) Scheme of the FR5 choice behavioral setup. Mice had free access to standard lab chow and could also lever press (FR5) for higher-palatable chocolate food pellets. (Right) Amount of food consumed represented as effort-based (FR5; solid) or freely-available lab chow (checkered). k, l. Break points for WT and NAc-D3RcKO mice during PR3 and PR7 sessions. Data in this figure and the rest of the manuscript are presented as mean ± SEM. Detailed figure statistics are included in Extended Data Table 1.

To further investigate whether cKO of NAc D3R expression impacted acquisition of motivation to run, we designed a choice task wherein mice were allowed to choose to spend time on either a fixed or a freely-moving disk, a distinct running apparatus, in an open field arena during the animal’s active cycle (Fig. 1e). WT-Cre and *Drd3*^fl/fl^-GFP control mice spent significantly more time on the freely-moving disk than on the fixed disk, in addition to showing robust running behavior (Fig. 1e-f). Conversely, NAc-D3RcKO mice did not significantly prefer the freely moving disk over the fixed disk and displayed minimal running (Fig. 1e-f). Running disks are angled relative to the ground (∼ 30° angle). As such, persistence is required for optimal running to develop since early attempts are marked by mice consistently falling off the disk. In control mice, preference for and running on the freely-moving disk did not begin immediately upon exposure but rather developed across the session as mice learned to run on the disk (Extended Data Fig. 1d). Consistent with increased persistence, control mice displayed increased entries into the freely-moving disk relative to NAc-D3RcKO mice (Extended Data Fig. 1e). These data provide support for NAc D3R in driving motivated running behavior, independent of diurnal cycle. We subsequently determined whether NAc D3R cKO would impact approach to other rewarding stimuli. We observed that NAc-D3RcKO mice displayed a non-significant decrease in sucrose preference relative to controls (Extended Data Fig. 1g). Further, this manipulation did not affect social approach (Extended Data Fig. 1h), suggesting that pursuit of low-effort palatable and social rewards is not affected by NAc D3R cKO. Anxiety-like behavior in the open-field and light-dark box (Extended Data Fig. 1f, i), as well novel object recognition (Extended Data Fig. 1j) was not different between controls and NAc-D3RcKO mice, demonstrating that exploratory behavior or interaction with novel stimuli are not impacted by NAc D3R cKO. These results collectively suggest that cKO of D3Rs in the NAc strongly disrupts running behavior associated with a motivated state.

We subsequently determined the role of NAc D3Rs in effort-related instrumental motivation, we used an operant conditioning paradigm in which animals had to lever press to acquire a chocolate pellet reward (Fig. 1g). Importantly, *Drd3* cKO in the NAc did not alter the body weight of NAc-D3RcKO mice relative to WT controls throughout the course of the experiment (Extended Data Fig. 2a). Animals first underwent an initial acquisition phase consisting of fixed-ratio (FR) 1 and 5 schedules to determine whether loss of NAc D3R signaling impacted reinforcement of food-seeking behavior. NAc-D3RcKO and WT groups displayed similar levels of active lever pressing and accuracy during FR1 sessions (Fig. 1h, Extended Data Fig. 2b-c), indicating that NAc D3R signaling is not essential for reinforcement. Interestingly, NAc-D3RcKO mice transiently displayed lower levels of responding when the effort required to obtain reward was increased from a FR1 to a FR5 schedule (Fig. 1i). To directly assess whether NAc D3Rs regulate motivation, mice subsequently underwent testing in an effort-based choice task in which they had the choice of consuming freely available standard chow or working for a higher-palatable chocolate reward^51,52^ (Fig. 1j, left). In this context, WT mice preferred to work for more palatable rewards over freely available lab chow (Fig. 1j). This pattern was absent in NAc-D3RcKO mice, which consumed higher quantities of regular chow than WT controls and did not show a preference towards working for a higher-palatable chocolate reward (Fig. 1j). Importantly, the overall amount of food consumed was similar across groups (Extended Data Fig. 2d), suggesting that there was a decrease in effort-based motivation but not homeostatic drive to eat in a hunger state. The same effect was observed when mice were given a choice between operant-obtained chocolate pellets and freely available chocolate pellets (Extended Data Fig. 2e-f). This indicates that control mice prefer to exert effort versus obtaining freely-available reward and that loss of NAc D3Rs biases choice towards reward-seeking behaviors that require less effort. Importantly, this is consistent with previous reports demonstrating that decreasing NAc DA signaling increases intake of low-effort food rewards and decrease effort or activity-based reward seeking behavior, without impacting overall food intake^51–56^. Subsequently, mice were subjected to progressive ratio (PR) schedules to further dissect the role of NAc D3R in regulating motivation. NAc-D3RcKO mice had decreased break points relative to WT controls in both PR3 (Fig. 1k) and PR7 schedules (Fig. 1l), as well as decreased PR session lengths (Extended Data Fig. 2g-h). These results indicate there is decreased motivation to exert effort to obtain food rewards with NAc D3R knockout. These data collectively indicate that motivated behavior necessary to obtain both appetitive food and running reward relies on NAc D3R function.

### NAc D3Rs are primarily expressed in D1-MSNs

The NAc is primarily composed of dichotomous cell types defined by DA receptor subtype (*i.e.*, D1R- and D2R-expressing MSNs), in addition to other molecular markers such as prodynorphin and proenkephalin^23^. To examine the NAc cell types in which D3R signaling occurs, we performed fluorescent *in-situ* hybridization experiments to detect the co-expression of *Drd1*a, *Drd2* and *Drd3* mRNA. *Drd3* mRNA, together with *Drd1a* and *Drd2*, was widely expressed in the NAc (Fig. 2a). Moreover, using *Drd3*-Cre mice crossed with tdTomato reporter mice, we observed robust tdTomato labeling in the NAc, in addition to the islands of Calleja (IC) (Extended Data Fig. 3a), but not the dorsal striatum. We found that in the NAc, a large majority (75.46% ± 0.89%) of *Drd3*-positive cells co-expressed *Drd1a*, with 20.51% ± 0.91% co-expressing *Drd2* (Fig. 2b-c). We also found that 81.79% ± 1.35% and 17.72% ± 1.32 of NAc *Drd1a*- and *Drd2*-positive neurons express *Drd3*, respectively (Fig. 2d), corroborating the finding that *Drd3*-expressing MSNs constitute a large subpopulation of D1-MSNs. Furthermore, the relative expression level of *Drd3* mRNA within the D1-MSN population was 20% higher than the level of *Drd3* expression in D2-MSNs (Fig. 2e). In addition, we obtained evidence of preferential expression of *Drd3* in NAc D1-MSNs when we recorded ChR2-mediated photocurrents in slices obtained from *Drd1a*-tdtomato/*Drd3*-Cre mice injected with AAV-DIO-ChR2-eYFP (Fig. 2f). Consistent with *in situ* hybridization results above, 77% of ChR2-positive neurons were positive for *Drd1a*-tdTomato (Fig. 2f). *Drd3* is therefore widely co-expressed in NAc D1-MSNs and less robustly in a subset of NAc D2-MSNs, suggesting that within the same neuronal population DA may serve different functions by acting on distinct receptor subtypes.

**Figure 2.**
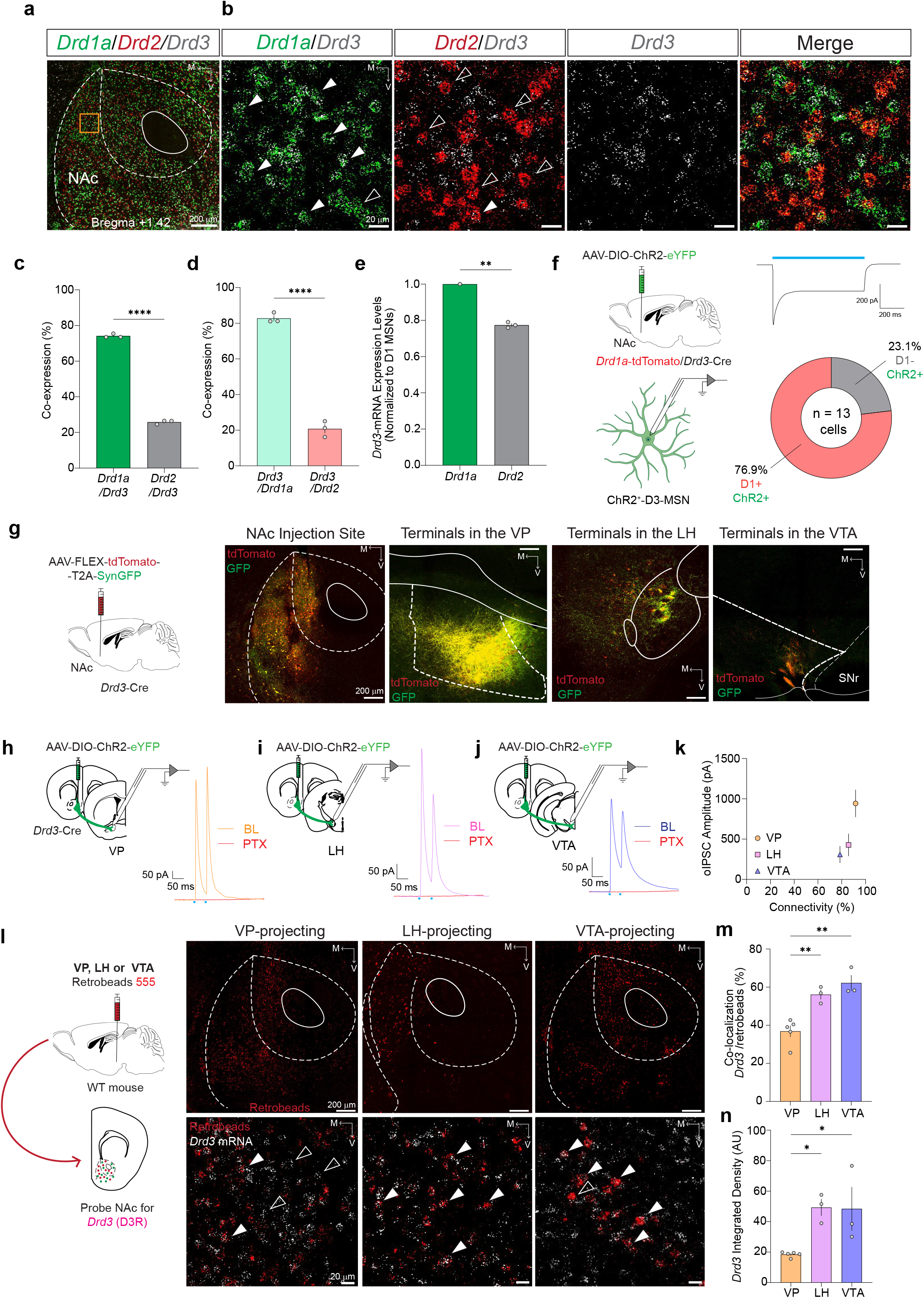
NAc D3Rs are primarily expressed in D1-MSNs and D3-MSNs display D1-MSN projection pattern (see also Extended Data Figure 3) a. Representative low-magnification confocal image of RNA *in situ* hybridization for *Drd1a* (green), *Drd2* (red), and *Drd3* (white) transcripts in the NAc. Orange inset shows the region targeted for zoomed images in b. b. Split high-magnification images of *Drd1a*/*Drd3*, *Drd2*/*Drd3*, and *Drd3* RNA expression in the NAc. Right image is an overlay of all channels. Filled arrowheads show co-labeled cells, and empty arrowheads show single-labeled cells. c. Percentage of *Drd3*^+^ cells co-expressing *Drd1a* or *Drd2* RNA in the NAc of WT mice. d. Percentage of *Drd1a*^+^ or *Drd2^+^* cells co-expressing *Drd3* mRNA in the NAc. e. Mean relative expression levels of *Drd3* mRNA (mean integrated density/area) relative to D1-MSNs in *Drd3*-mRNA-positive D2-MSNs. f. (Left) Schematic for quantification of *Drd3*^+^ cells using electrophysiological recordings. (Right) Representative trace showing light-evoked ChR2-mediated inward current in a ChR2-EYFP^+^ NAc MSN evoked by 1 s stimulation with 470 nm blue light. Pie-chart shows quantification of tdTomato-positive (*i.e.* D1-MSN) vs tdTomato-negative (putative D2-MSNs) in recorded ChR2 positive cells. g. (Left) Cre-dependent AAV-Syn-FLEX-tdTomato-T2A-SynaptophysinGFP was injected in the NAc of *Drd3*-Cre mice to visualize fibers (red) and synaptic terminals (green) in the outputs from D3-MSNs. (Right) Representative images showing a high density of NAc D3-MSN synaptic terminals in the VP, LH and VTA. h-k. (Right) Schematic of the electrophysiology experiment to assess functional connectivity from NAc D3-MSNs. A Cre-dependent AAV vector expressing ChR2-eYFP was injected in the NAc of *Drd3*-Cre mice. Acute slices containing the VP, LH or VTA were prepared from brains of *Drd3*-Cre mice 2–3 weeks after viral injection. (Left) Representative trace showing oIPSCs in VP, LH, and VTA cells. Red trace denotes bath-application of picrotoxin (PTX). k. Mean oIPSC amplitudes evoked by light stimulation of NAc D3-MSN to vs. connectivity of D3-MSNs to neurons VP, LH, and VTA. oIPSCs were detected in the majority of neurons recorded (VP, n= 11 of 12 neurons from 7 mice; LH, (n= 12 of 14 neurons from 8 mice; VTA, n= 11 of 14 neurons from 9 mice) l. Schematic of the retrograde tracing approach to compare NAc *Drd3*-expressing projection MSNs. Red retrobeads were injected into the VP, LH or VTA of WT mice. NAc sections were probed for *Drd3* mRNA) using *in situ* hybridization. Representative 20X confocal image showing retrobead labeling from VP-, LH-or VTA-projecting NAc MSNs (top). High-magnification images showing red-labeled retrobeads in the NAc co-localized with *Drd3* mRNA indicated by filled arrowheads (bottom). m. Quantification of the percentage of retrobead+ cells projecting to VP, LH or VTA that express *Drd3* in the NAc. n. Mean expression levels of *Drd3* mRNA (mean integrated density/area) in neurons projecting to VP, LH or VTA.

NAc MSNs play a role in the selection of appropriate goal-directed behaviors through connectivity with downstream brain regions. D1-MSNs project to the ventral pallidum (VP), lateral hypothalamus (LH) and ventral tegmental area (VTA), while NAc D2-MSNs projections are largely restricted to the VP^57–67^. To dissect the circuit-level mechanisms through which NAc D3R signaling-regulates motivated behavior, we determined the anatomical projections of *Drd3*-expressing MSNs using viral-assisted anterograde tracing in adult mice. Injection of AAV-FLEX-tdTomato-T2A-Synaptophysin-eGFP in the NAc of *Drd3*-Cre mice (Fig. 2g, left) allowed for the visualization of both fibers of passage and terminals of *Drd3*-expressing MSNs. We indeed observed putative presynaptic terminals labeled with Synaptophysin-eGFP puncta that were distinct from bundles of tdTomato-positive fibers, lacking eGFP (Supplementary Video 1). Consistent with the expression of D3Rs in D1-MSNs, GFP fluorescence revealed dense innervation within the NAc and of the VP, LH and VTA (Fig. 2g, right). Quantification of tdTomato fluorescence showed that fibers of passage were most prominent in more anterior regions (*i.e.*, VP), whereas putative terminals in the VP, LH or VTA (Extended Data Fig. 3b). Thus, D3-MSNs project downstream to the VP, LH and VTA and form discrete synaptic contacts in each of these regions.

We next used ChR2-assisted mapping to test the functional connectivity between NAc *Drd3*-expressing MSNs and their output regions. Patch clamp recordings in brain slices from *Drd3*-Cre mice injected with Cre-dependent ChR2 demonstrated optogenetically-evoked inhibitory post-synaptic currents (oIPSCs) recorded in the majority of VP, LH and VTA neurons (Fig. 2h-k). Light-evoked IPSCs were blocked by application of the GABA_A_-receptor antagonist picrotoxin (PTX) indicating that currents were mediated by GABA_A_-receptors (Fig. 2h-j). These findings indicate NAc D3-MSNs may regulate information processing in limbic circuits by sending inhibitory outputs to the VP, LH and VTA.

Distinct D1-MSNs have been shown to project to the VP, LH, and VTA^58,61–63^. Importantly these different MSN outputs do not collateralize, play distinct roles in behavior, and exhibit differential plasticity in response to behavioral experiences. To determine the proportion of D3R-expressing MSNs projecting to each of these three output regions, we performed retrograde tracing experiments in combination with *in-situ* hybridization (Fig. 2l). We found that, as reported, NAc MSN neurons projecting to these structures were anatomically segregated. VP-projecting MSNs were present in both NAc core and shell subregions (Fig. 2l, left). LH-projecting MSNs were primarily localized in the NAc shell (Fig. 2l, middle) whereas VTA-projecting MSNs were mainly located in the NAc core (Fig. 2l, right). In addition, quantification of cells positive for retrobeads and *Drd3* labeling showed that the proportion of D3R-expressing MSNs projecting to LH and VTA was higher than those projecting to VP (Fig. 2m). Further, *Drd3* mRNA expression was lower in VP-projecting MSNs, consistent with reduced expression in subsets of NAc D2-MSNs (Fig. 2n), which exclusively project to the VP. These results reveal the anatomical architecture by which *Drd3*-expressing MSNs connect within the NAc and to the VP, LH and VTA. These results led us to determine whether the role of NAc D3Rs in regulating motivation differed based on where the D3R expressing MSNs project.

### NAc D3R is essential for motivated behavior independent of projection neuron

NAc MSN projections to VP, LH and VTA have been previously shown to control reward-related behavior since distinct NAc projection neurons do not collateralize across these regions^58,59,62–68^, we hypothesized that D3R signaling in different MSN projections would differentially modulate motivated behavior. We tested this hypothesis by selective cKO of NAc *Drd3* expression in MSNs targeting either the VP, LH or VTA using intersectional viral and genetic approaches. To this end, we injected CAV-Flp-GFP in VP, LH or VTA of *Drd3*^fl/fl^ mice or WT control mice. This enabled expression of Flp recombinase in NAc MSNs targeting each of these distinct outputs (Fig. 3a). We then injected AAV-fDIO-Cre in the NAc to express Cre recombinase in retrogradely-infected NAc neurons expressing Flp and to effectively knockout D3Rs based on NAc outputs to either VP, LH, or VTA. In addition, we included AAV-FLEX-tdTomato into the NAc for histological confirmation of injection site (Fig. 3b, Extended Data Fig. 4). Three weeks after viral injection, mice were tested for motivation in running and operant tasks. Selective deletion of D3Rs from neurons projecting to either the VP, LH, or VTA in *Drd3*^fl/fl^ mice resulted in decreased motivation to run compared to WT controls, as reported in wheel running activity (Fig. 3c-e) and less preference for and running in the freely-moving disk in the disk choice task (Fig. 3f-h). Mice with cKO of D3Rs in either VP-, LH-, or VTA-projecting MSNs displayed diminished wheel running activity upon first exposure during the subjective day period (Fig. 3c-e), but not in response to diurnal shift to the animals’ inactive cycle (Extended Data Fig. 5a). The only exception were mice lacking D3R expression from VTA-projecting NAc MSNs, which displayed a decrease in dark cycle-initiated running (Extended Data Fig. 5a). Collectively, our findings indicate that D3R is essential for promoting motivated running behavior irrespective of their projections to downstream targets.

**Figure 3.**
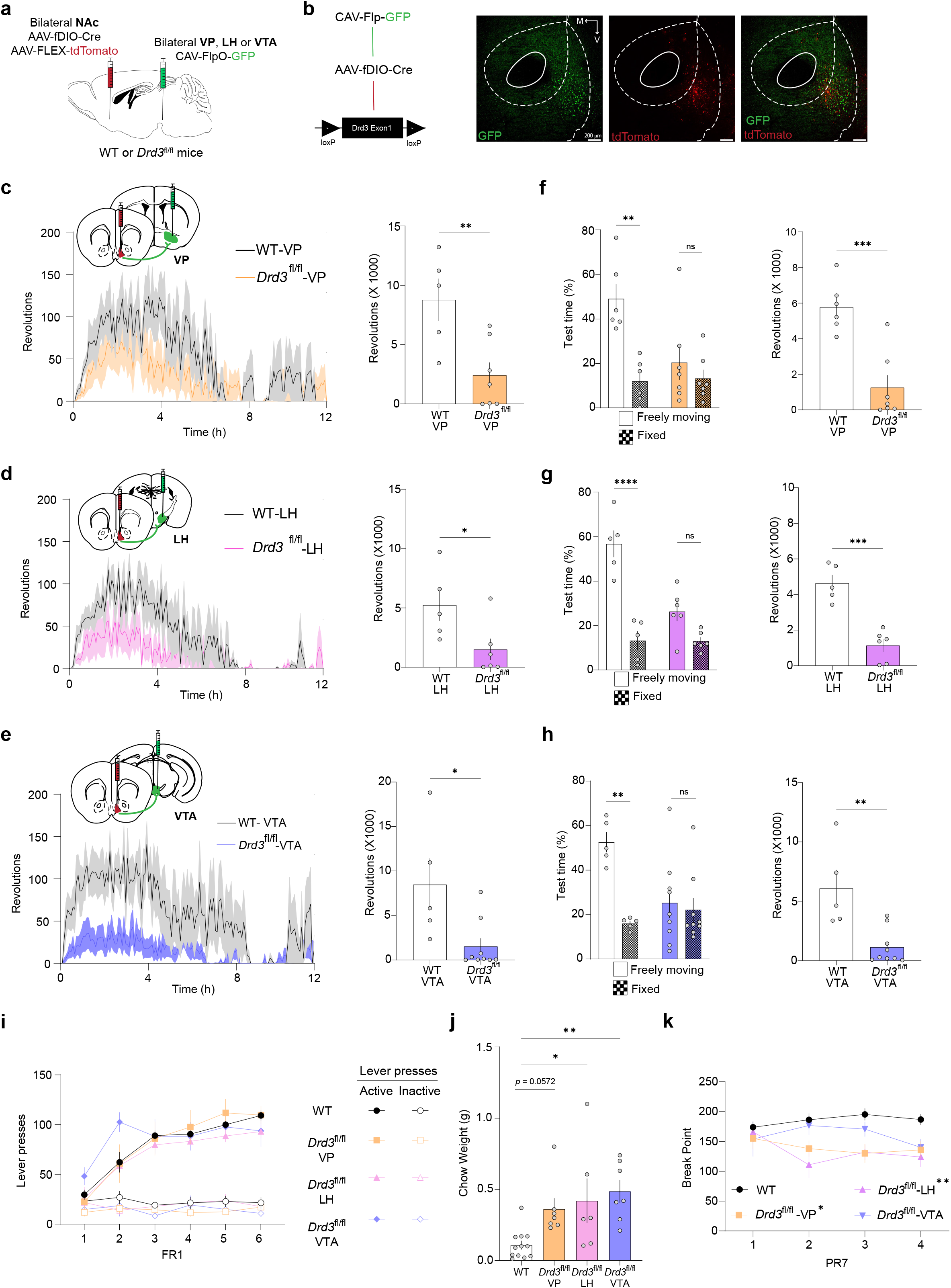
NAc D3R is essential for motivated behavior independent of projection neuron (see also Extended Data Figure 4 and 5) a. Diagram of viral injections for pathway-specific deletion of NAc *Drd3* from distinct MSN projections. Flp-dependent Cre and Cre-dependent tdTomato were injected bilaterally in the NAc of wild-type (WT) or *Drd3*^fl/fl^, and CAV-Flp-GFP was injected in the VP, LH or VTA to selectively knockdown *Drd3* expression in VP, LH or VTA-projecting NAc MSNs. b. (Left) Scheme showing fDIO-dependent Cre expression and recombination resulting in excision of exon 1 of the *Drd3* gene between flanking loxP sites. (Right) Representative images of GFP and tdTomato expression in NAc MSNs. Note: AAV-fDIO-Cre did not cause recombination in Ai14-tdTomato reporter mice injected in the NAc (Extended data Fig. 4). c-e. Time-course (left) and revolutions (right) of wheel-running activity during the first 12 hrs of running wheel exposure in WT or *Drd3*^fl/fl^ mice with pathway-specific deletion in the VP (Fig. 3c) LH (Fig. 3d) and VTA (Fig. 3e). f-h. Quantification of both time spent and wheel revolutions on the freely-moving and fixed disk for mice with the following injections: WT-VP and *Drd3*^fl/fl^ -VP (Fig. 3f), WT-LH and *Drd3*^fl/fl^ -LH (Fig. 3g), and WT-VTA and *Drd3*^fl/fl^ -VTA (Fig. 3h). i. Number of active and inactive lever presses of WT (black), *Drd3*^fl/fl^-VP (orange), *Drd3*^fl/fl^-LH (pink) and *Drd3*^fl/fl^-VTA (purple) animals during FR1 sessions. j. Amount of freely-available food consumed in the FR5 choice task. k. Break points for WT and *Drd3*^fl/fl^ mice during PR7 sessions.

We then assessed the effect of pathway-specific D3R cKO from either VP-, LH-, or VTA-projecting NAc MSNs in operant conditioning procedures. *Drd3*^fl/fl^ mice did not show changes in body weights (Extended Data Fig. 5b) or acquisition of responding on a FR1 schedule (Fig. 3i, Extended Data Fig 5c-d) relative to WT controls in all pathways. Interestingly, mice with selective cKO of *Drd3* from VTA-projecting NAc neurons had increased lever pressing under FR5 schedules. During choice tests, where animals could select between freely-available chow versus operant-derived chocolate pellets, selective cKO of D3Rs from either the VP-, LH-, or VTA-projecting MSNs increased intake of freely-available chow (Fig. 3j, Extended Data Fig. 5e-f). This was also observed when mice could choose between operant-derived and freely-available chocolate pellets (Extended Data Fig. 5g-i). These data indicate that *Drd3* cKO from either pathway biased consummatory behavior towards low effort rewards and away from higher effort rewards. NAc D3R cKO from VP- and LH-projecting MSNs resulted in deficits in motivation as shown by decreased breaking points in PR7 (Fig. 3k), but not PR3 (Extended Data Fig. 5j), consistent with decreased effort-based motivation. This effect did not reach statistical significance for mice lacking D3Rs in VTA-projecting MSNs. Our data collectively show that pathway-specific cKO D3Rs suppresses motivated behavior, but not reinforcement, independently of output region.

### D3R regulates GABAergic transmission from NAc collaterals and to the VP via a presynaptic site of action

G_i/o_-coupled GPCRs, such as D3Rs, regulate circuit function by inhibiting neurotransmitter release from axon terminals^69,70^. Local collaterals arising from NAc MSNs constitute a large proportion of inhibitory synapses onto MSNs, which are regulated by G_i/o_-coupled GPCRs and have been implicated in controlling striatal circuit recruitment during reward-related behavior^71–75^. To determine the functional role of D3R in shaping MSN function, we first examined the effect of D3R signaling on inhibitory synaptic transmission from MSN collaterals to neighboring MSNs. *Drd1a*-tdTomato/*Drd3*-Cre mice were injected with a virus expressing ChR2-eYFP in Cre-expressing neurons (Cre-ON; AAV-DIO-ChR2-eYFP) or Cre-negative neurons (Cre-OFF; AAV-DO-ChR2-eYFP), respectively, to selectively evoke GABA release from *Drd3*-positive or *Drd3*-negative NAc MSNs, respectively (Fig. 4a-b). Consistent with a higher proportion of *Drd3*-negative NAc neurons than *Drd3* positive, baseline oIPSC amplitudes were larger in the Cre-OFF condition (Extended Data Fig. 6a). Furthermore, in NAc MSNs recorded from mice expressing ChR2-eYFP in D3R-positive MSNs (Cre-ON), D3R activation with the D3R-selective agonist PD-128907 (1 µM) decreased oIPSC amplitude (Fig. 4c). Conversely, oIPSCs evoked from D3R-negative NAc cells (Cre-OFF) were insensitive to PD-128907 (Fig. 4c). These results suggest a presynaptic site of action for depression of D3-MSN collateral transmission by D3Rs. These results also validate the selectivity of PD-128907 for the D3R and selective viral-mediated transgene expression in D3R-expressing NAc MSNs in the *Drd3*-Cre mouse line. For confirmation, we also used the novel, highly selective D3R agonist ML417^76^, which also decreased oIPSC amplitude from NAc MSN collaterals (Extended Data Fig. 6b-d). In line with a role for presynaptic D3R in the regulation of NAc collaterals, the paired pulse ratio (PPR) increased (Fig. 4D), and the coefficient of variation (1/CV^2^) decreased (Fig. 4e) with PD-128907 in the Cre-ON condition. This reduction in GABA release probability was confirmed in experiments where the frequency of spontaneous inhibitory postsynaptic currents (sIPSCs), but not amplitude or kinetics, was decreased with PD-128907 application (Extended Data 6f-g), consistent with widespread D3R expression in D1-MSNs and a small subset of D2-MSNs. We next determined whether presynaptic D3R signaling differentially inhibited presynaptic D3-MSN collaterals onto D1- and D2-MSNs. First, D1-MSNs were identified by tdTomato fluorescence in *Drd1a*-tdTomato mice, while putative D2-MSNs were identified by lack of tdTomato fluorescence (Fig. 4b). Application of PD-128907 similarly inhibited oIPSCs evoked by optogenetic stimulation of D3-MSN collaterals in D1- and D2-MSNs (Extended Data Fig. 6h-i). Collectively, these results demonstrate that presynaptic D3R inhibits GABAergic outputs from MSN collaterals onto D1- and D2-MSNs.

**Figure 4.**
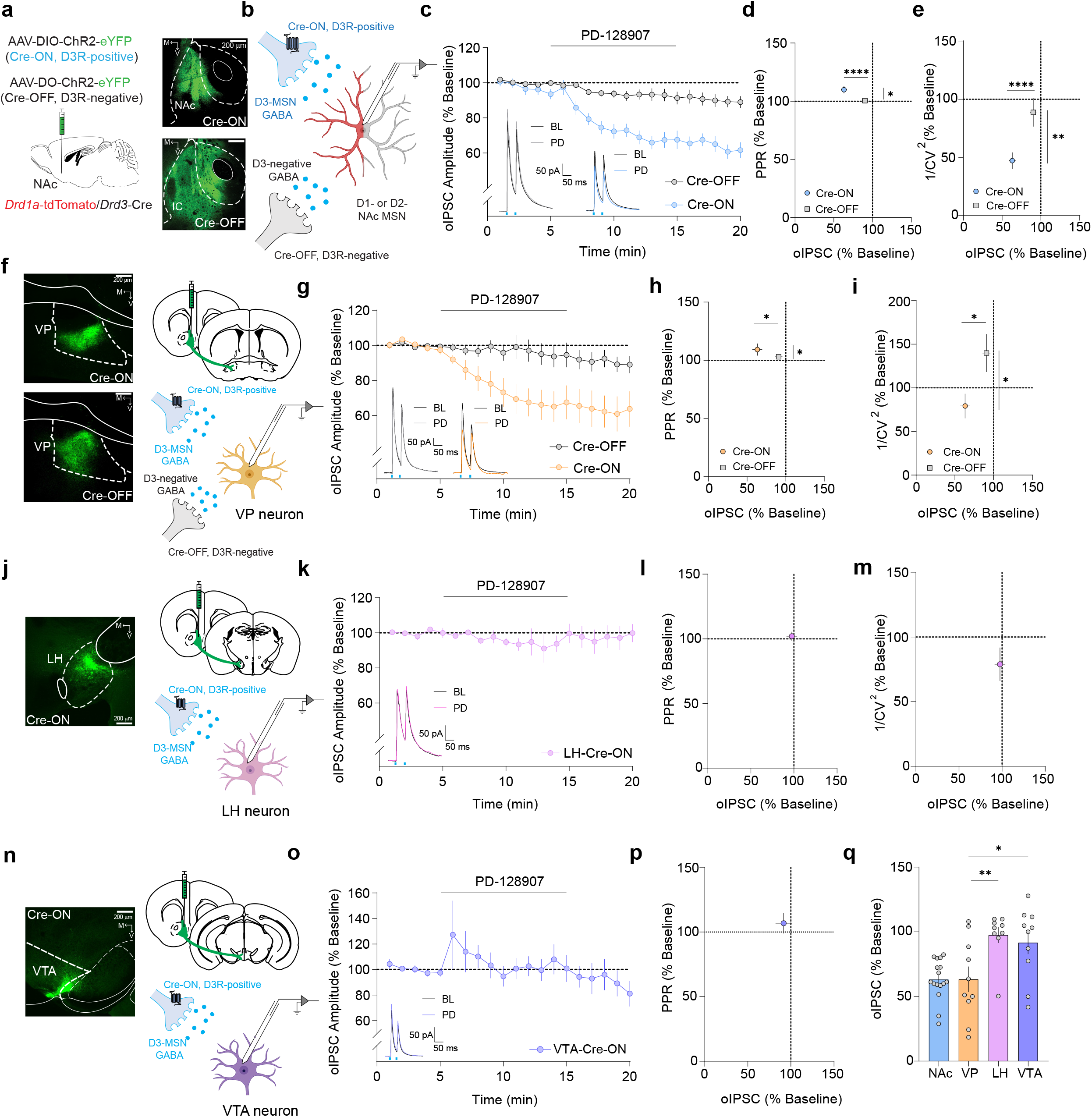
D3Rs regulate GABAergic transmission from NAc collaterals and to the VP via a presynaptic site of action (see also Extended Data Figure 6) a. (Left) Diagram of virus injection of AAV-EF1a-DIO-ChR2-eYFP (Cre-ON) and AAV-EF1a-DO-ChR2-eYFP (Cre-OFF) in the NAc of *Drd1a*-tdTomato/*Drd3*-Cre mice. (Right) ChR2-eYFP–expressing cell bodies of D3R-positive (top) and D3R-negative (bottom) terminals in the NAc. Note the lack of expression in the Islands of Calleja (IC) in mice expressing Cre-OFF ChR2 in the ventral striatum. b. oIPSCs originating from D3R-positive and D3R-negative collaterals were recorded from NAc D1- or D2-MSNs. c,g,k,o. Time-course of oIPSCs in NAc MSNs (Fig. 4c), VP cells (Fig. 4g), LH cells (Fig. 4k), and VTA cells (Fig. 4o) before, during and after bath application of the D3R-selective agonist PD-128907 (1 µM). For NAc MSNs and VP neurons Cre-ON groups are shown in blue (NAc MSN collaterals) or VP neurons (orange) while the Cre-OFF group is shown in black. (Inset) Representative oIPSCs traces recorded in NAc MSNs before and after bath application of PD-128907, d,h,l,p. Paired-pulse ratio (PPR, % baseline) versus oIPSC (% baseline) for NAc MSNs (Fig. 4d), VP cells (Fig. 4h), LH cells (Fig. 4l), and VTA cells (Fig. 4p). e,i,m. Coefficient of variation (1/CV^2^, % baseline) versus oIPSC (% baseline) for NAc MSNs (Fig. 4e), VP cells (Fig. 4i), and LH cells (Fig. 4m). f,j,n. (Left) Images of ChR2-eYFP-containing terminals in VP for Cre-ON (top) and Cre-OFF (bottom) conditions (Fig. 4f). Note the larger fiber density in the Cre-OFF condition arising from D3R-lacking D1-MSNs and D2-MSNs. (Right) oIPSCs originating from D3R-positive and D3R-negative MSNs were recorded from VP neurons. Fig. 4j and 4n show images of ChR2-positive fibers in the LH and VTA (left) and schematic depicting evoked GABA release from D3-MSNs onto LH and VTA cells, respectively (right). q. Summary graph of the inhibition of oIPSCs by PD-128907 from NAc, VP, LH or VTA neurons in Cre-ON condition.

Though NAc *Drd3*-expressing MSNs project to the VP, LH and VTA, it is unclear whether presynaptic D3Rs regulate GABAergic synaptic transmission from the NAc to each of these downstream targets. We therefore used ChR2-assisted functional mapping to examine the potential D3R modulation of synaptic connectivity between NAc D3-MSNs and VP, LH and VTA neurons. D3R activation using PD-128907 decreased GABA release onto VP neurons that was optogenetically evoked from *Drd3*-positive MSNs (Fig. 4f-g). This D3R-mediated decrease in GABA release was associated with an increase in PPR and decreased 1/CV^2^ (Fig. 4h-i). Conversely, GABA release from D3-negative MSNs targeted with Cre-OFF ChR2 onto VP neurons was not inhibited by PD-128907 (Fig. 4g). These findings demonstrate that D3Rs inhibit GABA release from MSNs onto VP neurons via a presynaptic site of action. Interestingly, D3R activation failed to decrease oIPSC amplitude in both LH and VTA neurons (Fig. 4j-q), suggesting that MSN connections onto LH and VTA neurons lack functional D3R. Taken together, our results reveal that presynaptic D3R on NAc MSNs selectively inhibits local collaterals and GABAergic transmission to the VP, but not in the outputs to the LH and VTA.

### Motivated behavior requires local D3R signaling within the NAc

Our electrophysiology results demonstrated that NAc collaterals and outputs to the VP, but not LH and VTA, are regulated by NAc D3Rs expressed in MSNs. However, cKO of D3Rs from NAc neurons projecting to VP, LH or VTA generally produced motivational deficits. Therefore, given that neurons projecting to VP, LH and VTA represent different populations, we hypothesized that D3R signaling is acting locally within the NAc to influence motivated behavior. We first tested whether microinfusion of the D3R antagonist ^77,78^ (1.8 ng per infusion site) into the NAc resulted in decreased running behavior (Fig. 5a). NAc D3R antagonism decreased preference for, and running on the freely-moving disk in the disk choice task (Fig. 5b), a result consistent with the hypothesis that D3R signaling within the NAc is essential for motivated running behavior. However, D3R antagonism within the NAc blocks D3R signaling in MSNs as well as any putative afferent inputs that may potentially express D3Rs at their terminals, such as the VTA, PVT or mPFC^79–81^. Thus, non-MSN D3Rs on afferent inputs might be contributing to D3R regulation of motivation. To specifically assess whether D3R acting on MSNs locally within the NAc microcircuitry is driving motivation, we implemented novel functional disconnection procedures involving pharmacological antagonism and cKO of NAc D3Rs (Fig. 5c). In these experiments, control *Drd3*^fl/fl^ mice received direct microinfusion of SB-277011A and ipsilateral injection of AAV-hSyn-GFP-Cre into the NAc (Fig. 5d). In this group, the unmanipulated hemisphere still has intact D3R signaling, hence supporting motivated behavior (Fig. 5e-f). Our experimental group (contralateral *Drd3*^fl/fl^ mice) underwent functional disconnections wherein unilateral NAc microinjections of SB-277011A and AAV-hSyn-GFP-Cre were made in contralateral hemispheres. Under these conditions, only D3R signaling in MSNs within the NAc underwent bilateral disruption (Fig. 5d). We posited that if motivated running behavior does not rely on local D3R signaling, then independently manipulating D3R activity in each of these hemispheres should not disrupt NAc D3R-mediated motivation. On the other hand, if motivation is mediated by local D3R signaling, then contralateral disconnection should disrupt motivation. Indeed, contralateral *Drd3*^fl/fl^ mice showed motivational deficits as shown by decreased running behavior and preference towards the freely-moving disk, (Fig. 5e-f) relative to their ipsilateral counterparts. As additional controls, we also quantified running disk performance in *Drd3*^fl/fl^ mice unilaterally injected with Cre-expressing virus or WT mice that had unilateral infusions of SB-277011A (Extended Data Fig. 7a-b). Consistent with the necessity of bilateral dysfunction in D3R signaling to observe changes in running choice behavior, unilateral disruption of D3R signaling by either cKO or pharmacological antagonism failed to modify preference for the freely-moving disk (Extended Data Fig. 7a-b). Taken together, these results suggest that D3Rs acting on MSNs locally within NAc microcircuitry is necessary for motivated behavior in mice.

**Figure 5.**
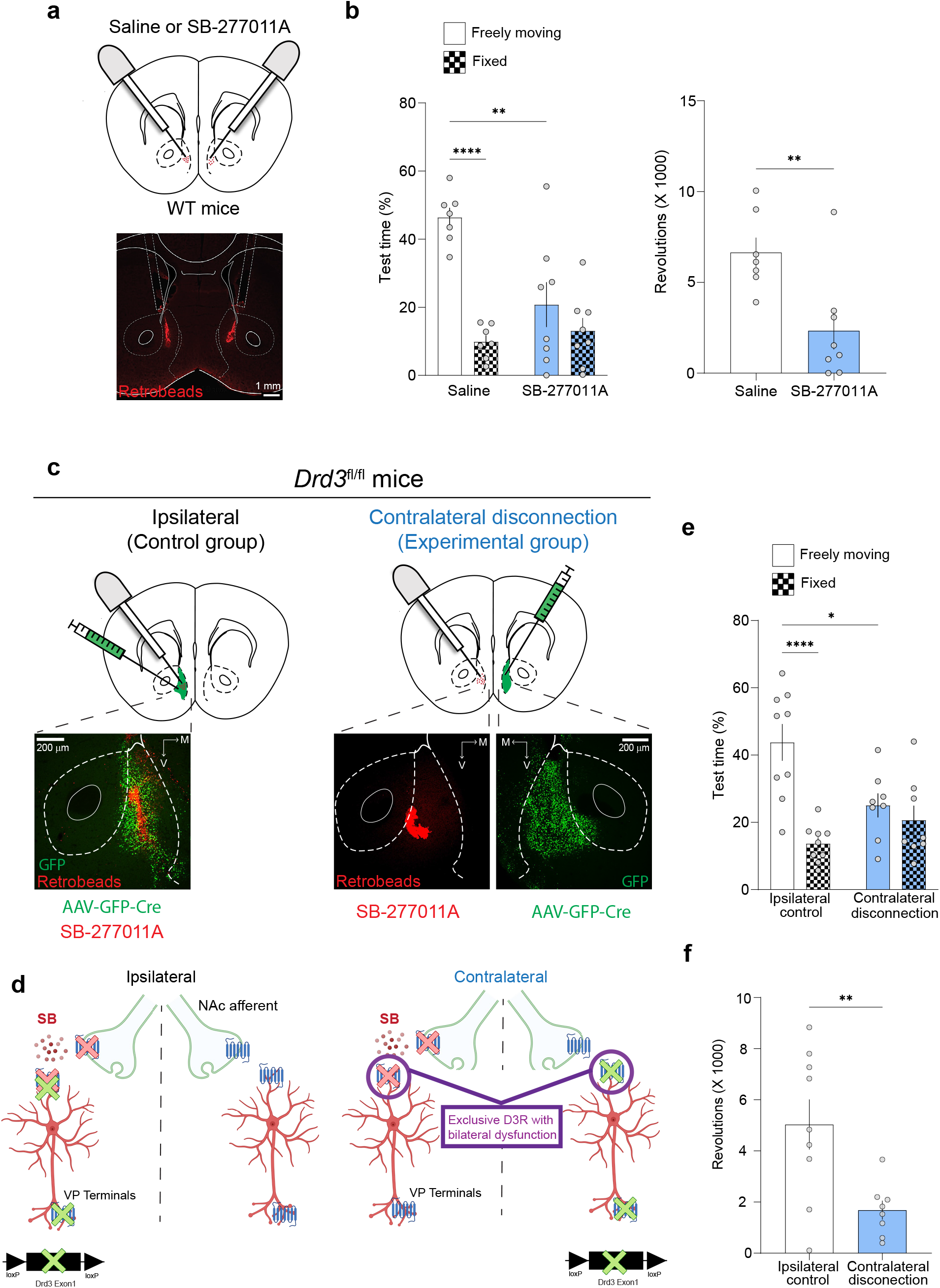
Motivated behavior requires local D3R signaling within the NAc (see also Extended Data Figure 7) a. Diagram (top) and representative image (bottom) of bilateral microinjection of the D3R antagonist SB-277011A (1.79 ng per hemisphere) into the NAc of WT mice. Red retrobeads were infused after the experiment to confirm accuracy of cannula placement. b. (Left) Percentage of time spent on the freely-moving and fixed disk for saline (white) and D3R antagonist (blue) groups during the running-disk choice task. (Right) Number of revolutions recorded in the freely moving disk. c. Diagram (top) and representative images (bottom) of selective inactivation of local NAc D3R function using functional disconnection experiments. Mice were injected unilaterally with AAV8-Syn-GFP-Cre into the NAc of *Drd3*^fl/fl^ mice. Ipsilateral (control) and contralateral disconnection (experimental) groups were infused with SB-277011A into the ipsilateral or contralateral NAc, respectively. d. Diagram describing rationale for disconnection procedures. In the ipsilateral group (left), one hemisphere was targeted with SB-277011A and AAV-GFP-Cre to suppress D3R signaling within the NAc and at terminals in the VP, while the other hemisphere had intact D3R signaling. For the contralateral group, the only common D3R dysfunction in both hemispheres was local D3R signaling within the NAc (purple circles), which was targeted by antagonist infusion and contralateral injection of AAV-GFP-Cre. e. Percentage of time spent on the freely-moving (solid) and fixed disk (checkered) for ipsilateral control (white) and contralateral disconnection (blue) groups. f. Number of revolutions registered in the freely moving disk.

Specific effects of D3R signaling on MSN physiology, such as inhibition of local collaterals and outputs to the VP, but not LH and VTA, may underlie regulation of motivation. We therefore determined the effects of non-selectively activating G_i/o_-coupled GPCRs using chemogenetics throughout the cell to determine how this differed from endogenous D3Rs which have location-specific effects on MSNs and drive running. To this end, we bilaterally injected AAV-hSyn-DIO-hM4D(G_i_)-mCherry or AAV-hSyn-DIO-mCherry in the NAc of *Drd3*-Cre mice and subjected the mice to the running disks choice task (Extended Data Fig. 7c). Chemogenetic inhibition with clozapine N-oxide (CNO) 30 min prior to the start of testing significantly reduced preference for the freely moving disk and decreased running behavior in mice expressing hM4Di relative to mCherry controls (Extended Data Fig. 7d). hM4D-mediated suppression of NAc D3-MSN activity did not affect, however, general locomotor activity or open field anxiety (Extended Data Fig. 7e). Of note, this approach causes DREADD overexpression in all compartments of D3-MSNs ^82^, including dendrites and outputs to LH and VTA, where we showed NAc D3R do not regulate synaptic transmission. This further emphasizes that the specific presynaptic distribution of D3Rs in NAc MSNs is essential to drive motivational running. Thus, taken together, our results suggest that D3R-induced decrease in GABAergic output drives motivation to run via presynaptic inhibition of NAc MSN collaterals.

### NAc D1Rs mediate reinforcement, but not motivation

Since *Drd3* is largely co-expressed with *Drd1a* D1Rs and the vast majority of D1-MSNs express *Drd3* (Fig 1), we determined if these DA receptor subtypes differ in controlling reward function. As low-affinity receptors, NAc D1Rs modulate reward-related behaviors presumably by detecting phasic changes in DA and modifying synaptic transmission onto MSNs^25–27^ However, the precise role of NAc D1Rs, as compared to D3Rs, in regulating motivation and reinforcement remains unclear. *Drd1a*^fl/fl^ mice injected with AAV-Cre (NAc-D1RcKO) into the NAc were run concomitantly with the NAc-D3RcKO experiments described above (Fig. 6a). We found that cKO of *Drd1a* in the NAc resulted in impaired FR1 acquisition, suggesting that D1R signaling in the NAc is necessary for reward-driven reinforcement (Fig. 6b). The decrease in lever pressing in NAc-D1RcKO mice persisted under FR5 schedules of reinforcement and dissipated with training (Fig. 6c). Similar to WT controls and opposite NAc-D3RcKO mice, NAc-D1RcKO mice preferred to work for a higher palatable reward in the effort-based choice task (Fig. 6d). Furthermore, break points during progressive ratio schedules did not differ between WT and NAc-D1RcKO mice (Fig. 6e), indicating that the motivation to seek a palatable reward remained intact. Similar to the lack of effect in instrumental motivation, NAc *Drd1a* cKO did not strongly impact motivated running behavior in the running wheel and disk choice behaviors (Fig. 6f-g, Extended Data Fig. 8a). Furthermore, NAc *Drd1a* cKO did not have alter sucrose preference (Extended Data Fig. 8b), sociability (Extended Data Fig. 8c), light-dark box (Extended Data Fig. 8d) or novel object recognition (Extended Data Fig. 8e) tests, indicating that NAc D1R function is not essential for hedonic processing, anxiety-like and novelty-seeking behaviors, respectively. Taken together, our results suggest that D1Rs, but not D3Rs, play a role in reinforcement.

**Figure 6:**
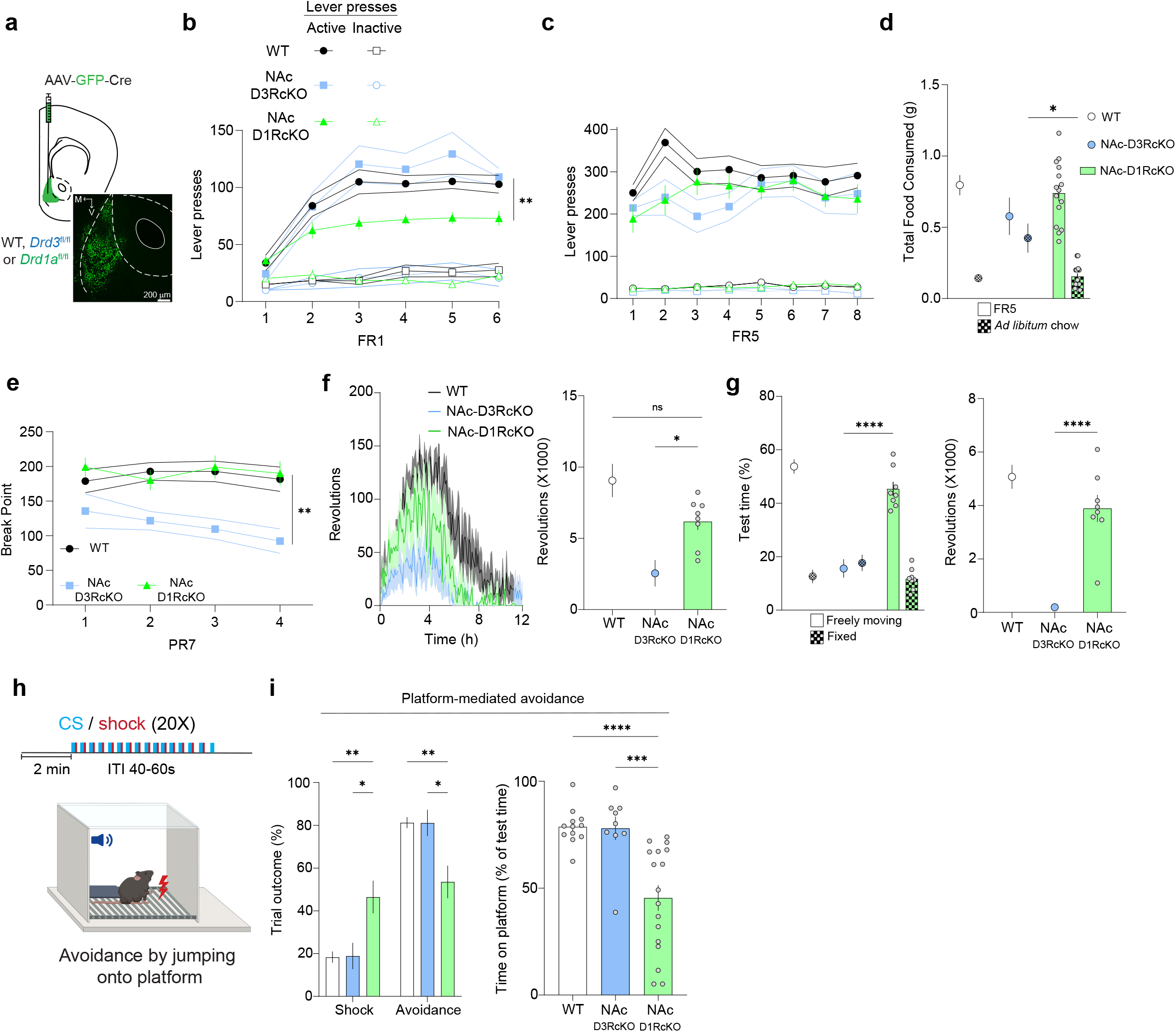
NAc D1Rs mediate reinforcement but not motivation (see also Extended Data Figure 8) a. Scheme (top) and representative image (bottom) depicting the NAc area targeted with AAV-GFP-Cre (green) for experiments shown in b-i. b-c. Number of active (filled) and inactive (unfilled) lever presses of WT (black), NAc-D3RcKO (blue) and NAc-D3RcKO (green) animals during sessions with FR1 and FR5 schedules of reinforcement. d. Amount of food consumed during the effort-related choice task represented as FR5 effort-based (solid) or freely-available lab chow (checkered). e. Break points for WT, NAc-D3RcKO and NAc-D1RcKO mice during PR7 sessions. f. (Left) Timecourse of wheel-running activity during the initial 12 hr period of wheel exposure (in the animal’s inactive cycle) in WT-Cre, NAc-D3RcKO and NAc-D1RcKO mice. (Right) Quantification of revolutions during the 12 hr period. g. Quantification of time and revolutions for WT, NAc-D3RcKO and NAc-D1RcKO groups during the running disk choice task. (Left) Percentage of test time spent on the freely-moving and fixed disk. (Right) Number of revolutions recorded in the freely moving disk for each group. h. Scheme of the platform-mediated avoidance task. Mice were placed in an operant box and were presented with 20 pairings of conditioned stimulus (CS) and a footshock as the unconditioned stimulus (US). Animals could step onto the platform to actively avoid a footshock. i. Quantification of trial outcome (avoidance or shock) as percentage of total trials for day 1. j. Percent time spent on the platform. Note: operant and running data from WT and NAc-D3RcKO groups were acquired concomitantly with NAc-D1RcKO mice and were replicated from Fig. 1. Thus, data from WT and NAc-D3RcKO groups are displayed differently (mean and error bars).

The role of NAc D1Rs may not be limited to positive reinforcement, but may also play a role in reinforcement of threat avoidance (negative reinforcement) in addition to positive reinforcement. We determined whether reinforcement of avoidance of a footshock requires NAc D1Rs using a modified platform-mediated avoidance paradigm^83,84^. Here, mice were presented with an auditory cue (CS, tone) that co-terminated with a footshock (Fig. 6h), and mice could avoid the footshock by mounting a platform located in one of the corners of the chamber. NAc-D1RcKO had avoidance deficits and received more shocks and spent significantly less time on the platform than WT mice (Fig 6i). In contrast, NAc-D3RcKO mice did not display deficits in shock avoidance. Deficits in avoidance behavior in NAc-D1RcKO mice were also observed upon re-exposure to the platform-mediated avoidance task 24 hours later (Extended Data Fig. 8f), supporting a sustained deficit in threat avoidance. These results further demonstrate that NAc D1R, but not D3R, activity is essential for negative reinforcement, suggesting that D1Rs regulate both positive and negative reinforcement. Collectively, these results show that D3Rs and D1Rs play dissociable roles in regulating motivation and reinforcement, respectively.

### D3R and D1R regulate separable features of NAc D1-MSN physiology

We hypothesized that the dissociable roles of D1Rs and D3Rs in motivation and reinforcement may be a consequence of divergent physiological effects in NAc D1-MSNs. Previous studies have proposed that D1R regulation of N-methyl-D-aspartate receptor (NMDAR)-dependent plasticity is a fundamental mechanism driving reward learning^26,27,85^. Using glutamate uncaging to evoke isolated postsynaptic NMDAR currents (Fig. 7a), we found that activation of NAc D1Rs with the D1-like receptor agonist SKF-81297 potentiated AP-5-sensitive NMDAR currents in D1R-positive neurons, an effect that was blocked with bath-application of the D1R-selective antagonist SCH39166 (Fig. 7b-d). Interestingly, D3R activation failed to modify NMDAR currents (Fig. 7b-d). Furthermore, SKF-81297 failed to potentiate NMDAR currents in D1R-positive projection neurons with D1R cKO, demonstrating the specificity of our pharmacological and genetic manipulations (Fig. 7e-h). Thus, D1Rs, but not D3Rs, potentiate NAc NMDAR currents, which provide a potential substrate for reward learning and reinforcement. We subsequently demonstrated that D1R activation using SKF-81297 did not modify GABA release from D3R MSNs NAc collaterals (Fig. 7i-l), consistent with separable control of inputs and outputs of NAc MSNs by D1Rs and D3Rs, respectively. Altogether, these results demonstrate that dissociable control of motivated and reinforced behaviors by D1Rs and D3Rs likely stems from dissociable physiological effects in D1-MSNs.

**Figure 7:**
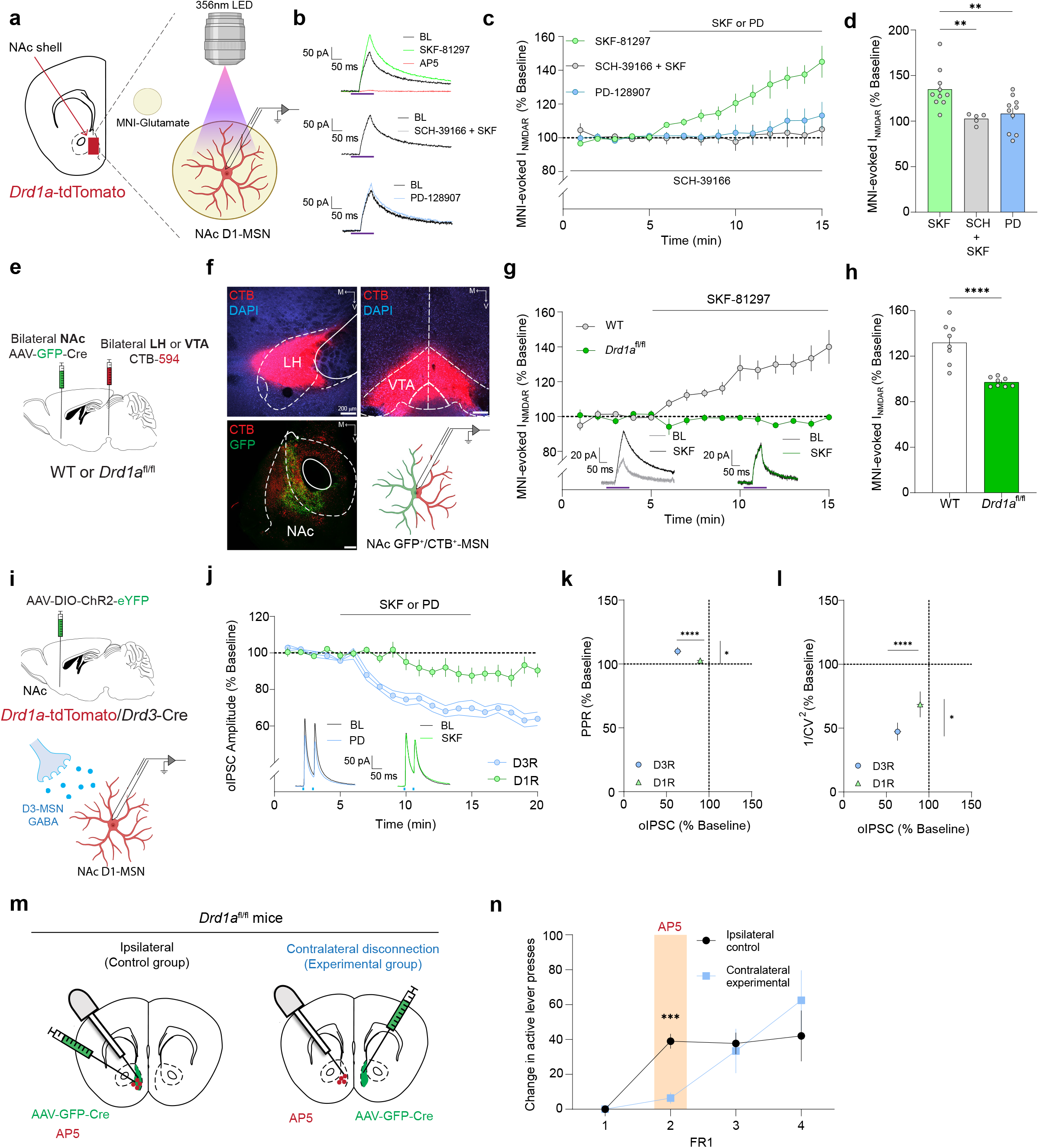
D3R and D1Rs regulate separable synaptic features of NAc D1-MSNs (see also Extended Data Figure 8) a. Schematic showing location of patch-clamp recordings (left) and glutamate uncaging (right). MNI-Glutamate (50 µM) was uncaged using 365 nm UV light (150 ms pulses) and biophysically-isolated NMDAR-currents were recorded at +40 mV from NAc D1-MSNs using DNQX (10 µM), PTX (50 µM) and tetrodotoxin (TTX; 1 µM). b. Representative traces of evoked NMDAR currents in NAc D1-MSNs before and after bath-application of the D1R agonist SKF-81297 (green, top), application of SKF-81297 in the presence of the D1R antagonist SCH-39166 (grey, middle) and application of the D3R agonist PD-128907 (blue, bottom). Red trace indicates that evoked NMDAR currents were eliminated with AP5 (50 µM). Purple bars indicate ultraviolet light pulses. c. Time-course of normalized NMDAR-current amplitude in NAc D1-MSNs before and during bath application of SKF-81297 (10 µM, green), preincubation with SCH-39166 (1 µM) and application of SKF-81297 (black) or application of PD-128907 (1 µM, blue). d. Evoked NMDAR currents after treatment with SKF-81297, preincubation with SCH-39166 and treatment with SKF and application of PD-128907 (% baseline). e. Schematic of experimental details to demonstrate D1R-NMDAR interactions and validation of *Drd1a* cKO in NAc of *Drd1a*^fl/fl^ mice. AAV8-Syn-GFP-Cre was injected bilaterally in the NAc of WT or *Drd1a*^fl/fl^ mice to genetically ablate *Drd1a*, and the retrograde tracer CTB-594 was injected in the LH or VTA to selectively label NAc D1-MSNs. f. Representative images of CTB injection sites in LH and VTA, and colabelling of GFP and CTB in the NAc. Recordings were made from colabeled GFP^+^/CTB^+^ NAc MSNs. g. Time-course of normalized NMDAR-current amplitude in NAc D1-MSNs before and during and after bath application of SKF-81297 in WT or *Drd1a*^fl/fl^ groups. (Inset) Representative NMDAR traces recorded in NAc D1-MSNs for each genotype. h. I_NMDAR_ (% baseline) change after treatment with SKF-81297 for WT and *Drd1a* groups. i. Schematics of experiment to determine regulation of NAc collaterals by D1Rs and D3Rs. *Drd1a*-tdTomato/*Drd3*-Cre mice we injected with AAV5-Ef1a-DIO-ChR2-eYFP (Cre-ON) in the NAc (top) and GABA release was evoked from D3R-positive terminals (1ms pulses). Biophysically-isolated oIPSCs were recorded from NAc D1-MSNs (bottom). j. Time-course of oIPSCs in NAc MSNs from D3R collaterals before, during and after bath application of SKF-12897 or PD-128907. (Inset) Representative traces recorded in NAc MSNs before and after bath application of PD-128907 (blue) or SKF-81297 (green). Note: data using PD-12897 were replicated from Fig. 4 and was therefore displayed with different error bars. k. Paired-pulse ratio (PPR, % baseline) versus oIPSC (% baseline). l. Coefficient of variation (1/CV^2^, % baseline) versus oIPSC (% baseline). m. Diagram of disconnection procedures of D1R and NMDAR function in the NAc. Mice were injected unilaterally with AAV-Syn-GFP-Cre into the NAc of *Drd1a*^fl/fl^ mice. Ipsilateral control (left) and contralateral disconnection (right) groups were infused with AP5 into the ipsilateral or contralateral NAc, respectively. n. Change in active lever presses lever presses under an FR1 schedule of reinforcement relative to Day 1 in ipsilateral (black) and contralateral (blue) groups. AP5 was microinjected on Day 2 of FR1 sessions.

Lastly, to determine whether D1R interactions with NMDAR currents are necessary for reinforcement, we performed disconnection experiments of NAc D1R and NMDAR function during acquisition of FR1 lever pressing. *Drd1a*^fl/fl^ mice were injected with AAV-GFP-Cre and infused with AP5 (700 ng per infusion site) in the ipsilateral hemisphere (control group) or injected with AAV-Cre and infused with AP5 in the contralateral hemisphere (experimental group) (Fig. 7m). AP5 microinjection on day two of FR1 acquisition session decreased FR1 lever pressing (Fig. 7n), consistent with deficits in reward reinforcement. Importantly, decreased lever pressing was not influenced by the number of lever presses during the Day 1 baseline session (Extended Data Fig. 8g). Decreased FR1 lever pressing was not sustained since levels of lever pressing were not significantly difference the day after AP5 microinjection. Collectively, our results reveal key role of NAc D1Rs in the expression of positive and negative action reinforcement. Mechanistically, this process involves, at least in part, amplification of postsynaptic NMDAR-mediated excitatory drive onto NAc D1-MSNs.

## Discussion

Here, we have identified distinct roles of NAc D3Rs and D1Rs in regulating different features of D1-MSN physiology that promote complementary features of reward-related behaviors. Activation of presynaptic D3Rs, but not D1Rs, in NAc MSNs inhibits GABAergic transmission from collaterals within local microcircuits to promote effort-based motivation. Conversely, NAc D1Rs, but not D3Rs, regulate reinforcement by interacting with NMDARs. These findings indicate that the effects of DA on NAc MSNs are more complex than canonical models of striatal function where D1Rs and D2Rs are largely segregated, and provide a new model wherein DA receptor signaling at distinct receptors (D1R and D3R) within a single cell type can differentially mediate motivation and reinforcement.

Our integrative approach of genetic and pharmacological manipulations of D3R expression and function enabled us to isolate the roles of NAc D3R signaling in reward function, surmounting challenges associated with earlier approaches that primarily relied on pharmacology (*i.e*., lack of cellular or receptor specificity) and constitutive knockouts (*i.e*., lack of regional/cellular specificity and possible developmental compensations). We employed two different paradigms that permit assessment of motivation, instrumental and running behaviors. We showed that NAc D3R cKO disrupts motivation as assessed by decreased break points in PR schedules in operant settings, a standard measure of motivation. Further, NAc D3R cKO biased choice for low-effort rewards versus high-effort rewards, consistent with a loss in motivation. Since food-seeking driven by hunger was intact in NAc D3R cKO mice, but effort-based motivation was decreased, this suggest that NAc D3Rs regulate properties of motivation that rely on efforts but not homeostatic drive. Importantly, in other striatal compartments, such as the dorsal striatum, D3Rs are not as enriched and the role this region plays in motivation is less established^31^. Here, we also utilized running, a multi-dimensional behavior, to further demonstrate that NAc D3Rs regulate the motivational component of a non-food based rewarding stimulus. We found that NAc D3R signaling was necessary for promoting running on a novel wheel during their inactive cycle and in a choice task during the active cycle, but not general locomotor activity or wheel running under the control of circadian cycle. Indeed, DA signaling in the NAc has been recently shown to underlie motivation to exercise^86^. Together, results from operant and running procedures are consistent with the hypothesis that D3Rs regulate the activational components of motivation, which include effort, persistence, and/or vigor^3,8,11,12,87^. This is of particular importance since physical activity and effort, or lack thereof, has been linked to fluctuations in motivation in various neuropsychiatric diseases^88^.

Our study suggests that DA acting presynaptically on D3Rs inhibits GABAergic inhibitory transmission arising from local MSN collaterals to drive motivated behavior. D3R suppression of local inhibitory connections might disinhibit MSNs and concomitantly activate downstream targets of the NAc to promote motivation. Consequently, this work advances our previous understanding of the behavioral relevance of NAc collateral connections, which have been posited to regulate striatal-dependent behavior^71,75^. Furthermore, NAc D1-MSNs also corelease neuropeptides, such as dynorphin and substance P^9^, which mediate reward and avoidance behaviors when acting in the dorsomedial or ventromedial NAc shell^89^. Thus, D3Rs may orchestrate multimodal signaling from MSNs by regulating local collateral outputs. Additionally, D3Rs may regulate different aspects of D1-MSN physiology, such as intrinsic excitability^37^, within the NAc. Nevertheless, whether these physiological actions of D3R signaling are also important for regulating reward function is presently unclear.

Coexpression of *Drd3* and *Drd1a* in NAc D1-MSNs might provide these cells with the ability to detect DA in different spatiotemporal domains. Striatal MSNs, and NAc MSNs by extension, have been classically defined by the expression of a single DA receptor subtype^23,24^, and this is thought to underlie differential effects on reward function. In this study, we have shown that the majority of NAc D1-MSNs highly express *Drd3*, and only a subset of D2-MSNs express *Drd3*. Furthermore, *Drd3* is expressed at higher levels in D1-MSNs than D2-MSNs, suggesting D3R is anatomically poised to complement D1R function in D1-MSNs. We demonstrated that D3Rs and D1Rs are likely providing NAc D1-MSNs with different computational properties via divergent cellular effects in distinct sub-cellular compartments. Electron microscopy studies have shown that D1Rs and NMDARs colocalize in the somatodendritic compartment of NAc MSNs^90^. This is consistent with the present and previous electrophysiological findings showing that D1Rs potentiate NMDA receptor currents or D1R-NMDA receptor interactions are critical for plasticity^26,27,85,91–93^. Further, we found that D3R signaling inhibits GABA release from MSN collaterals, providing evidence that D3Rs are functionally localized to MSN axon terminals. This dissociation suggests that functional D1R and D3R may be localized to distinct sub-cellular compartments. D1Rs and D3Rs may also both regulate common aspects of MSN physiology not explored in the present study. Even under those circumstances, D3Rs and D1Rs are coupled to inhibitory G_i_ and stimulatory G_s_ proteins, respectively^35^, and these differences in coupling to downstream G protein effectors could also promote the differential effects on MSN physiology in domains where the two receptors may overlap. Further, D3Rs have 100-fold greater affinity for DA than D1Rs^18,35^, which is likely to influence how these receptors are engaged by complex DA dynamics. As lower affinity receptors, D1Rs may only be occupied by large increases DA concentrations associated with phasic DA release. This provides a mechanism to potentiate NMDAR activity at excitatory synapses that were coincidentally active during phasic DA release and subsequently promote long-lasting plasticity, such as long-term potentiation mediated by structural changes and increased AMPAR currents^26,27^. In contrast, D3Rs may be occupied by “tonic” DA signaling generated by spontaneous DA transients or be useful for detecting dips in DA transmission. Further, in contrast to D1Rs, D3Rs would not only be occupied during the peak of phasic DA release but also during the decay phase of the phasic DA response. Together, unique co-expression of D1R and D3R provides NAc D1-MSNs with the molecular machinery to orchestrate dissociable features of reward-related behavior via differential translation of distinct modes of DA transmission into physiological changes.

To our knowledge, this is the first demonstration that separate DA receptors provide co-expressing NAc MSNs exert distinct physiological effects to regulate separate features of reward-related behavior. Since D3Rs and D1Rs are present within the same D1-MSN population and exert different cellular effects and have different affinities for DA, this work resolves long-standing questions concerning what behaviors DA signals control at different local concentrations and timescales. Within this framework, we predict that D3Rs would detect slower changes in DA (*i.e.*, changes in tonic DA driven by internal state), and regulate motivation through inhibition of NAc collateral transmission. Conversely, we found that D1Rs acting on postsynaptic NMDAR-mediated currents are critical for reinforcement of reward-seeking behavior. NAc D1R may be necessary for the early stages of learning when reward- and punishment-evoked transients are largest and cue evoked-transients are emerging. Thus, phasic DA acting through NAc D1Rs would permit D1 MSNs to integrate specific representations carried by coincidentally-active excitatory afferent inputs via NMDA-dependent plasticity and changes in excitability^94^. This mechanism would provide D1 MSNs the capacity to strengthen representations conveyed by upstream excitatory neuronal ensembles converging on individual D1-MSNs to support execution of reinforced behaviors. Motivation and reinforcement are dissociable reward features with complementary roles in establishing and maintaining goal-directed behavior, and coordinated D3R and D1R signaling may help integrate these functions. For example, as DA unbinds D1Rs (lower affinity DA receptor) during the decay phase of phasic DA release triggered by reinforced behavior, D3Rs (which binds DA with higher affinity) may remain bound and activated and increase motivation in response to feedback from a positively reinforced outcome. This is an important consideration given that DA kinetics in the ventral striatum are longer-lasting relative to the dorsal striatum^95^. Our model therefore provides a receptor-based mechanism by which motivational states can be adjusted by on-going behavior depending on outcomes. Lastly, DA release can be regulated locally in the NAc independently of VTA DA neuron firing, including via cholinergic and opioid receptors that directly excite and inhibit DA terminals, respectively^96–101^. This is of relevance since DA release controlled at the level of the VTA or locally may differentially contribute to DA dynamics and influence how D1Rs and D3Rs in D1-MSNs are activated.

In conclusion, our work refines our circuit-level understanding on how DA release in the NAc is translated into dissociable aspects of reward function via D3Rs and D1Rs co-expressed in NAc MSNs. This provides a novel description of separable control of reward and physiological features by DA within the same cell type. A plethora of psychiatric and neurological disorders are characterized by dysfunctional or amplified reinforcement and/or motivation. The present study provides insight into how dysfunction in DA release and/or signaling in brain disorders may impact specific reward domains via NAc D1Rs and D3Rs, and provides potential therapeutic targets to treat specific alterations in distinct reward features.

## Methods

### Mice

Adult female and male mice (aged 8-20 weeks at the start of experiments) were used throughout the study. No significant differences were found between both sexes, and data were therefore pooled to complete final group sizes. DA D3 receptor-IRES-Cre (*Drd3*-Cre) (Tg(Drd3-cre)KI196Gsat/Mmucd, GENSAT, KI196, a gift from Charles Gerfen) and *Drd3*-Cre crossed with Ai14-tdTomato reporter mice (*Drd3*-Cre/Ai14) were made congenic with a C57BL/6J background and were used for anatomical characterization and electrophysiological experiments. *Drd3*^fl/fl^ mice (kindly provided by Z. Freyberg) were also bred on a C57/BL6J background and were used for behavioral experiments.

Generation of *Drd3*^fl/fl^ mice: To develop this strain, LoxP elements were inserted flanking the transcriptional start site in exon 1 of *Drd3* (Extended Data Fig. 1a). Specifically, a targeting vector was designed via recombineering as described previously ^102,103^. We first retrieved approximately 12.2 kb of *Drd3* genomic sequence encompassing 8.3 kb of the 5’-upstream region preceding exon 1 through 3.5 kb of the intron 1 sequence from the BAC, RP24-135K7. This genomic sequence was inserted into a pDTA vector containing the PGK-DTA negative selectable marker by gap repair. We then inserted the 5’ LoxP site approximately 4 kb upstream of the *Drd3* transcriptional start site in exon 1 followed by insertion of Frt-PGFneo-Frt-LoxP approximately 500 bp 3’ downstream of exon 1. The final vector contains 5’ and 3’ arms of 4.2 kb and 3.2 kb, respectively. The vector was then linearized by NotI digestion, purified and electroporated into mouse ES cells derived from an F1(129Sv/C57BL/6J) blastocyst. Electroporated cells were cultured in the presence of G418 48-hrs post-electroporation to select for cells with successful genomic integration of our construct. G418-resistant colonies were subsequently picked and screened by long range PCR using primers corresponding to sequences outside the arms and specific to the 5’ and 3’ LoxP sites to identify targeted ES clones. Targeted ES clones were then expanded and further analyzed by long-range PCR for confirmation prior to using them for ES-morula aggregation to generate chimeric animals. The resulting chimeric mice were then bred with ROSA26-FlpoER mice to remove the PGKneo cassette to generate the final *Drd3*^fl/fl^ mice. These *Drd3*^fl/fl^ mice were then made congenic with the C57BL/6J genetic background via backcrossing for 10 generations (N10). For PCR genotyping, the following primers were used: *Drd3* Lox gtF 5′-TGAGACTAAGCAGCGTCCAC-3′, *Drd3* Lox gtR 5′-CTCTGAGTTAGATCTCCCCAGC-3′ for WT 372bps/Floxed 468 bp and *Drd3* Frt gtF 5′-GCTGGCTCTCCATAGATTCTGC-3′, *Drd3* Frt gtR 5′-CTTGAACAGATGTAGGCACCCTG −3′ for WT 254 bp/Floxed 347 bp.

*Drd1a*^fl/fl^ mice were acquired from The Jackson Laboratory (JAX, 025700) and were used for behavioral and electrophysiological experiments. *Drd1a*-tdTomato (B6.Cg-Tg(*Drd1a*-tdTomato)6Calak/J, JAX, 016204) and *Drd1a*-tdTomato/*Drd3*-Cre mice were used for electrophysiological experiments. Wild-type (WT; C57BL/6J, JAX#000664) mice were also used in all experiments and were bred at NIMH or obtained from Jackson Laboratories. *Drd3*-Cre mice used for anatomical and electrophysiology experiments were heterozygous.

Mice were group housed (2–5 mice per cage) in temperature- (21-24 °C) and humidity- (40-65%) controlled facilities and maintained on a reverse 12-h light/12-h dark cycle with lights off at 8 am. All mice were maintained in filter-topped cages and provided food and water *ad libitum*, except for animals undergoing testing in operant procedures. Single housing was necessary for experiments requiring food restriction (operant conditioning) or acclimation to behavioral tasks (wheel running experiments) and is explicitly denoted in those cases. All purchased mice were kept in the local animal facility for at least one week following delivery before initiating experimental procedures. Mice were monitored for health status daily and before experimentation for the entirety of the study. All efforts were made to minimize pain and distress and the number of mice used. All procedures were performed in accordance with the National Institutes of Health (NIH) Guide for the Care and Use of Laboratory Animals and approved by the Animal Care and Use Committee of the National Institute of Mental Health Intramural Research Program.

### Viral constructs

Recombinant adeno-associated viruses (AAVs) and type 2 canine adenoviruses (CAV2) were implemented to allow expression of transgenes of interest. AAVs were purchased from Addgene, the NIDA Genetic Engineering and Viral Vector Core, Boston Children’s Viral Vector Core and UNC Viral Vector Core. CAV2 was acquired from the Institut de Génétique Moléculaire de Montpellier. All constructs were aliquoted and stored at −80 °C. Titers ranged from 10^12^ to 10^14^ genomic copies per mL. Specific details on each viral construct can be found in Extended Data Table 2.

### Stereotaxic surgeries and optical fiber / guide cannula implantation

All surgeries were conducted under aseptic conditions and body temperature was maintained at approximately 36 °C with a heating pad. Mice, 8-16 weeks of age, were anaesthetized with a mixture of ketamine (100 mg/kg body weight; intraperitoneal injection) and xylazine (10 mg/kg body weight; intraperitoneal injection) as confirmed by complete absence of flinching response to pinch. The animal’s head was shaved, and ophthalmic ointment (GenTeal) was applied to the eyes to prevent drying. Mice were subsequently placed in a stereotaxic apparatus (David Kopf Instruments Model 1900, Tujunga, CA, USA) and the surgical site was exposed using a sterile scalpel after cleaning with povidone–iodine and 70% ethanol. The mouse’s head was leveled by ensuring the difference in dorsoventral distance between bregma and lambda was within 50 µm. A small craniotomy window was then made above the injection site with a stereotax-mounted drill. The following injection coordinates (in mm) and volumes were used: [anterior-posterior (AP) and medial-lateral (ML) relative to bregma; dorsal-ventral (DV) relative to dura mater at target coordinate]: NAc (AP: +1.40; ML: ±1.65; DV −4.50, 12° angle towards midline; 300 nL), VP (AP: +0.40; ML: ±1.35; DV −5.35; 300 nL), LH (AP: - 1.35; ML: ±1.10; DV −5.25; 300 nL), VTA (AP: −3.30; ML: ±1.85; DV −4.60, 14° angle; 300 nL). Infusions were made at a rate of 100 nL/min utilizing 29 Ga microinjection needles connected to FEP tubing secured to a 2 µl Hamilton syringe and a microinjection pump (UMP3, World Precision Instruments, Sarasota, FL). The infusion system was filled with distilled water and separated from the infused virus or drug by a small air bubble. The injector tip was first lowered 100 µm deeper than the target DV coordinate and then raised to the planned coordinate before infusion to facilitate viral diffusion at the site of injection, instead of along the needle track. After infusion, the injector was kept for 8 min at the injection site to allow for diffusion and was then slowly withdrawn.

For cKO of *Drd1a* or *Drd3* in the NAc, homozygous *Drd1a*^fl/fl^ or *Drd3*^fl/fl^ mice received bilateral injections of AAV8-hSyn-GFP-Cre (4.50E+12 GC/mL) or AAV1-EF1α-eGFP in the NAc; WT mice were used as controls.

For anatomical tracing experiments, *Drd3*-Cre mice were injected with AAV2/9-phSyn1(S)-FLEX-tdTomato-T2A-SynaptophysinEGFP or AAV1-Syn-FLEX-Chrimsom-tdTomato (4.10E+12 genome copies (GC)/mL) into the NAc to characterize downstream projections and terminals of NAc D3R-containing neurons.

For pathway-specific deletion of NAc *Drd3*, CAV-Flp-GFP (1.05e13 GC/mL) was delivered to the VP, LH or VTA and a mixture of AAV5-EF1α-fDIO-Cre (5.00E+12 GC/mL) and AAV1-CAG-FLEX-tdTomato (5.90E+12 GC/mL) was delivered to the NAc at a ratio of 9 to 1, respectively.

For electrophysiological studies examining D3-MSN connectivity and D3R modulation of synaptic transmission, *Drd1a*-tdTomato/*Drd3*-Cre mice received bilateral injections of AAV5-EF1α-DIO-ChR2(H134R)-eYFP (4.00E+12 GC/mL) or AAV1-EF1α-DO-hChR2(H134R)-eYFP (5.88E+12 GC/mL) targeting the NAc.

For drug microinjection experiments, mice were bilaterally implanted with stainless steel guide cannulas (26-gauge, 3.5 mm in length, P1 Technologies) 1 mm above the NAc (AP: +1.40; ML: ±1.65; DV −3.50 from dura, 12° angle towards midline). Guide cannulas were then secured to the skull using MetaBond cement and dummy cannulas were used to maintain cannula patency and removed only during the injection period.

For disconnection procedures, *Drd3*^fl/fl^ or *Drd1a*^fl/fl^ mice were injected with AAV8-Syn-GFP-Cre and implanted with a guide cannula above the NAc in the same hemisphere (ipsilateral group) or injected with virus and implanted with the guide cannula in the contralateral hemisphere (contralateral group).

For validation of the *Drd1a*^fl/fl^ line, *Drd1a*^fl/fl^ mice received bilateral injections of Alexa Fluor 594-conjugated Cholera Toxin Subunit B in PBS (CTB-594, 1.0 mg mL^−1^, no. C-34776, Thermo Fisher) in either LH or VTA brain regions.

For chemogenetic inhibition experiments, *Drd3*-Cre mice received bilateral injections of AAV1-EF1α-DIO-hM4D(Gi)-mCherry in the NAc to inhibit D3R-expressing NAc MSNs. AAV5-hSyn-DIO-mCherry (1.2E+13 GC/mL) was used for control groups.

Following all surgical procedures, incisions were closed using VetBond (3M, Maplewood, MN) or surgical staples. Mice were allowed to recover from anesthesia in heating pads until they showed regular breathing and locomotion, at which point they were transferred back to the vivarium. Animals received subcutaneous injections of ketoprofen (5 mg/kg body weight) for three consecutive days for post-operative analgesia and anti-inflammatory purposes. Experiments involving the use of AAVs were initiated 3-4 weeks after injection, 3 weeks for CAV2-Flp-GFP and 7 days for retrobead and CTB injection procedures.

### Anatomical characterization of *Drd3*-expressing MSNs

#### RNAscope fluorescent *in situ* hybridization

Multiplex fluorescence *in situ* hybridization (RNAscope, Advanced Cell Diagnostics, Newark, CA) was used to detect the expression of *Drd1a*, *Drd2*, *Drd3*, and *Cre* mRNAs in the NAc of adult WT and *Drd3*^fl/fl^ mice. For all RNAscope procedures, tools, slides, and equipment were cleaned with 70% ethanol and RNAse inhibitors (RNAZap, Invitrogen) prior to use. Mice were euthanized by cervical dislocation, brains were rapidly dissected, and flash-frozen for 20 seconds in 50 mL of 2-methylbutane chilled on dry ice. Subsequently, brains were stored at −80°C until sectioning. 16-µm slices containing the NAc were obtained using a cryostat (CM3050 S; Leica Biosystems; Deer Park, IL, USA) at −20 °C and thaw-mounted onto Superfrost microscope slides (Fischer Scientific) in a series of four slides. Slides containing sections were stored at −80°C until *in situ* hybridization processing. *Drd1a*, *Drd2*, *Drd3* or *Cre* mRNA signal was detected using the RNAscope fluorescent kit following ACDBio manual^104^. Briefly, slides containing the NAc were removed from −80°C, fixed with prechilled 4% paraformaldehyde for 20 min at 4°C, and subsequently washed twice for 1 min with PBS, before gradual dehydration with 50% ethanol (1 x 5 min), 70% ethanol (1 x 5 min), and 100% ethanol (2 x 5 min). Next, slides were air-dried at room temperature for 10 min and a hydrophobic barrier was drawn around the slides using a hydrophobic pen (Vector Laboratories, Newark, CA). Sections were then incubated with Protease Pretreat-IV solution for 20 min at room temperature. Slides were washed with ddH2O (2 x 1 min), before being incubated with the appropriate probes for 2 hr at 40°C in the HybEZ oven (Advanced Cell Diagnostics) and undergoing hybridization. Probes used were purchased from Advanced Cell Diagnostics as follows: Mm-*Drd1a*-C1 (nucleotide target region 444-1358; Accession number NM_010076.3), Mm-*Drd2*-C2 (nucleotide target region 69-1175; Accession number NM_010077.2), Mm-*Drd3*-C3 (nucleotide target region 23-1253; Accession number NM_007877.1), *Cre* recombinase (nucleotide target region 2-972; Accession number N/A). Probes were warmed-up to 40°C in a water bath until use.

Slides were washed in wash buffer twice for 2 min, prior to being incubated with three drops of amplification 1 buffer, Amplification 2 buffer, Amplification 3 buffer and Amplification 4-Alt A/C buffer at 40°C in the HybEZ oven for 30, 15, 30, 15 minutes respectively. Slides were washed in wash buffer twice for 2 min. DAPI solution was applied to sections at RT for 20 sec to label nuclei. Finally, slides were coverslipped using Vectashield Hard Set mounting medium (Vector Laboratories). Slides were stored at 4°C until imaging. Z-stacked images including the NAc were acquired using an A1R confocal microscope (Nikon, Tokyo, Japan) with a 20X objective. This produced a tiled image containing the entirety of the NAc which was used for quantification. The following combinations of laser excitation and emission filters were used for various fluorophores: DAPI (405 nm excitation; 450/30 nm emission), eGFP (491 nm laser excitation; 528/38 nm emission), tdTomato (561 nm laser excitation; 586/15 nm emission), Cy5 (647 nm laser excitation; 665/35 nm emission). All samples were imaged with the same settings to allow comparison between samples. Background subtraction and thresholds were set uniformly for all images.

#### Quantification of *in-situ* hybridization

Image analysis and cell quantification were performed using ImageJ software (Fiji, version 2017) and CellProfiler software (version 3.1.9.; Broad Institute; Cambridge, MA) ^105,106^. To analyze the images, each image was opened in ImageJ and converted to a maximum intensity projection. Images were overlapped onto the Allen Mouse Bran atlas to set boundaries for the total NAc area to be analyzed. Two to three serial sections (between approximately AP +1.42 and +1.21) were analyzed on the total NAc area for each mouse. For quantification, images were imported to an automated CellProfiler pipeline that was kept identical across samples from the same experiment. Here, only cells with a clear DAPI+ nucleus were counted, which were then registered and used as markers for individual cells. ROIs for analysis were defined as the 3 µm area surrounding the DAPI signal. A blinded experimenter set thresholds for each channel which determines the minimum intensity of fluorescence for a probe to be counted. These thresholds were validated by visual spot check throughout the image to ensure cells and probes were being appropriately counted. CellProfiler software provided CSV files with the total counts of cells and levels of overlap, which are reported in the data. For co-expression of *Drd1a*, *Drd2* and *Drd3*, cells considered as positive consisted of an area within the radius of a DAPI nuclear staining that measured at least 10 positive pixels for receptor probes. For the percentage of retrobead-positive cells expressing *Drd3* mRNA, retrobead-positive cells contained at least 6 for retrobead labeling.

#### Anterograde tracing of D3R-expressing NAc MSNs

To examine the projection pattern of *Drd3*-expressing NAc neurons, 300 nL of AAV1-hSyn1-FLEX-Chrimsom-tdTomato or AAV2/9-phSyn1(S)-FLEX-tdTomato-T2A-SynaptophysinEGFP-WPRE were bilaterally injected into the NAc of *Drd3*-Cre mice as described above. Mice were perfused three weeks following viral injection and 50-μm-thick coronal brain slices were prepared. Images were taken at approximately 300-μm intervals from brain regions expressing tdTomato and/or GFP using a Nikon A1R confocal microscope with a 20X objective. Regions of interest were labeled relative to bregma based on the “Paxinos and Franklin’s The Mouse Brain in Stereotaxic Coordinates” (Franklin, K. B. J. & Paxinos; Academic Press, an imprint of Elsevier, 2013). Total integrated fluorescence intensity tdTomato and GFP from each downstream target was quantified using ImageJ with identical pinhole, gain, and laser settings. For each brain region, four images were acquired at the same focal point from each animal. No additional post-processing was performed for any of the images analyzed here.

#### Retrobead retrograde tracing of D3R-expressing NAc MSNs

C57BL/6J WT mice were injected bilaterally with 200 nL of Red and Green Retrobeads IX (Lumafluor, Durham, NC) into the VP, LH or VTA/SNc as described above. Per the manufacture’s protocol, red Retrobeads were injected at a 1:4 dilution and green Retrobeads were left undiluted. Seven days after injection, brains were collected and processed for RNAscope procedures as described above.

### Quantitative real-time PCR

WT and *Drd3*^fl/fl^ mice expressing Cre-recombinase in the NAc were euthanized by cervical dislocation. Brains were rapidly dissected, and 1 mm coronal brain slices containing the ventral striatum were obtained by slicing the brain placed in an iron matrix (Kent Scientific, Corp., Torrington, CT). The NAc was microdissected bilaterally and immediately transferred to microcentrifuge tubes on dry ice; samples were stored at −80°C for RNA isolation and processing. Total RNA was extracted from dissected NAc samples using the NZY RNA Total Isolation kit (ref. MB13402, Nzytech, Lisboa, Portugal), and purified mRNA samples were reverse transcribed using the SuperScript IV First-strand cDNA synthesis kit (Thermo Fisher Scientific). Target sequences were amplified from the cDNA using the TaqMan Gene Expression Assay Kit (Thermo Fisher Scientific) and the SYBER Green system (Power SYBR Green PCR Master Mix, Applied Biosystems). All Taqman probes were purchased from Applied Biosystems and were as follows: *Drd1a* (Mm02620146_s1), *Drd2* (Mm00438545_m1), *Drd3* (Mm00432887_m1), GAPDH (Mm99999915_g1). Quantitative PCR (qPCR) was performed using TaqMan Fast Polymerase (Applied Biosystems, Waltham, MA) in an ABI PRISM 7900HT SDS Real-Time PCR system ((Applied Biosystems). Cycling conditions were as follows: initial hold at 95°C for 20 s; 40 cycles of step 1 (95°C for 1 s); and step 2 (60°C for 20 s). Samples were run in triplicates, and negative controls were run in parallel. The relative mRNA expression level for each sample was calculated using the ΔΔCt method, where Ct was the cycle threshold for each reaction and GAPDH was used as internal control (DCt = Ct (gene of interest) – Ct (GAPDH)^107^. Gene expression fold change was calculated by normalizing the value of each sample to the mean of the control samples.

### Histology

Upon completion of all experiments, mice were deeply anaesthetized with euthanasia solution (VedCo Inc., St. Joseph, MO), and then transcardially perfused with 40 mL of cold phosphate-buffered saline (PBS 1X, pH 7.4), followed by 40 mL of cold 4% w/v paraformaldehyde (PFA) in PBS. Brains were extracted and post-fixed in 4% PFA at 4 °C overnight and then cryoprotected in 20% sucrose-PBS for 24 hours, followed by 30% sucrose-PBS for 24-hrs, at which point they were stored in PBS or prepared for sectioning on a cryostat. To this end, brains were embedded on the mounting disk with Tissue-Tek Optimum Cutting Temperature Compound (Sakura Finetek USA, Torrance, CA) for freezing over dry ice. Brains were subsequently placed in the cryostat at −20 °C, and consequently sectioned into 50- or 100-μm slices. Slices were mounted on slide glasses with DAPI Fluoromount-G mounting medium (0100-20, Southern Biotech, Homewood, AL) for visualization on a Nikon A1R confocal microscope (10X objective, NA 0.45, lasers: 405 nm, 488 nm, 561 nm). Injection sites and optical fiber placements were routinely confirmed in all animals by preparing coronal sections containing the region of interest. After histological verification, animals with insufficient transgene expression, off-target transgene expression outside the region of interest by visual inspection, and/or inaccurate implant placement were excluded from data analyses. A representative scheme of the viral spread for NAc-D3RcKO mice included in this study is included in Extended Data Fig. 1k.

### *Ex-vivo* electrophysiology

Slice electrophysiology recordings were performed as previously described^54,75^. Briefly, 3 to 8 weeks after surgery, mice were deeply anaesthetized with euthanasia (200mg/kg ip; VedCo, Inc.) and subsequently decapitated after confirmation of absent toe and tail pain reflexes. Brains were rapidly removed and chilled for 2 min in ice-cold NMDG-based slicing solution containing (in mM): 92 NMDG, 20 HEPES, 25 glucose, 30 NaHCO_3_, 2.5 KCl, 1.2 NaH_2_PO_4_, 5 sodium ascorbate, 3 sodium pyruvate, 2 thiourea, 10 MgSO_4_, and 0.5 CaCl_2_ (pH 7.35, 303-306 mOsm) and saturated with 95% O_2_/5% CO_2_. Brains were rapidly blocked, dried on filter paper, and glued to a platform containing ice-cold NMDG slicing solution within a vibratome (VT1200, Leica). Coronal sections (300 µm) containing the NAc, VP, LH and VTA were cut at a speed of 0.07 mm/s while the brain was submerged in ice-cold NMDG-based slicing solution. Following slicing, sections were kept in a custom-built chamber containing NMDG slicing solution for 7 min at 34°C. Slices were subsequently transferred to a chamber filled with modified holding aCSF saturated with 95% O_2_/5% CO_2_ containing (in mM): 92 NaCl, 20 HEPES, 25 glucose, 30 NaHCO_3_, 2.5 KCl, 1.2 NaPO_4_, 5 sodium ascorbate, 3 sodium pyruvate, 2 thiourea, 10 MgSO_4_, and 0.5 CaCl_2_ (pH 7.35, 303-306 mOsm) at room temperature for at least 1 hr. Slices remained in this solution for recovery until transfer to the recording chamber. For recordings, the recording chamber was kept at 31°C and perfused with a pump (World Precision Instruments) at a flow rate of 1.5-2.0 mL per minute with aCSF containing (in mM): 126 NaCl, 2.5 KCl, 1.4 NaH_2_PO_4_, 1.2 MgCl_2_, 2.4 CaCl_2_, 25 NaHCO_3_, and 11 glucose (303-305 mOsm) at 31°C. Cells were visualized with a 40X water-immersion objective on an Olympus BX5iWI inverted microscope equipped with infrared-differential interference contrast (IR-DIC) optics and epifluorescence (Olympus Corp, Tokyo, Japan). 4IPatch pipettes (2-4 MΩ) were pulled from borosilicate glass (G150TF-4; Warner Instruments, Hamden, CT) and filled with a freshly filtered (0.22 µm syringe filter) cesium-based internal solution (in mM: 117 cesium methanesulfonate, 20 HEPES, 0.4 EGTA, 2.8 NaCl, 5 TEA-Cl, 4 Mg-ATP, 0.4 Na-GTP and 5 QX-314 (pH 7.35; 280-285 mOsm). Whole cell access was obtained from individual neurons after acquisition of a giga-ohm seal recording. All recordings were made utilizing a Multiclamp 7400B amplifier (Molecular Devices) and data were digitized at 10 kHz and filtered at 1-2 kHz using a 1440A Digidata Digitizer (Molecular Devices). Series and input resistances (10-20 MΩ) were monitored online using a −5 mV, 70-ms voltage pulse through the recording electrode. Cells with >20% change in access resistance were discarded from further analysis. Liquid junction potentials were ≈ −7 mV and were left uncorrected. Data were analyzed offline using Clampfit 10.6 (Molecular Devices, San Jose, CA).

To isolate GABA_A_ responses evoked by optogenetic stimulation of D3R-expressing MSNs, the AMPA receptor antagonist DNQX (10 µM) and NMDAR antagonist D-AP5 (50 µM) were included in the aCSF, and neurons were voltage-clamped at 0 mV. Optogenetic-evoked inhibitory postsynaptic currents (oIPSCs) were elicited every 10 s by photostimulating ChR2 using two 1-ms pulses of 473-nm LED light (pE-300^ultra^, CoolLED, Andover, United Kingdom) separated by a 50-ms interstimulus interval. ChR2-negative cells were identified by the lack of ChR2 currents evoked by blue light stimulation. ChR2 currents were characterized by sustained, steady state currents in response to 1-s blue light stimulation with an onset immediately at the start of the laser pulse. Synaptic GABAergic currents in ChR2-negative cells were outward-currents that were not sustained for the duration of a 100-1000 ms blue light pulse and had a delayed onset beyond the 1 ms optical pulse. Cells that did not show a peak that exceeded baseline noise in this window were counted as non-responders. Connectivity was calculated as the percentage of cells receiving oIPSCs from NAc D3R-expressing MSNs. Paired pulse ratios were calculated as the amplitude of the second peak divided by the amplitude of the first peak. IPSCs were recorded until their amplitudes were stable for at least 5 minutes, at which point PD-128907 (1 µM), ML417 (1 µM) or SKF 81297 (10 µM) were added to the bath for 10 min. After bath application, drugs were washed out for 5 additional min. Timecourse graphs of the effects of drug application were generated by averaging raw oIPSCs measurements in 1-min bins and expressing each point as a percentage of the average of the 5-min baseline. The averaged baseline and 5-min after drug application were used for quantification of modulation of synaptic transmission. To differentiate between D1- and D2-MSNs, slices were obtained from *Drd1a*-tdTomato mice and recording were made from tdTomato positive (i.e. D1-MSNs) or tdTomato-negative (putative D2-MSNs) cells. Because the electrophysiological results from the two MSN populations were similar, data were grouped. Recordings in the VP, LH and VTA were made irrespective of the identity of the cell. Spontaneous IPSCs (sIPSCs) were collected in the presence of DNQX (10 µM) and D-AP5 (50 µM) and the last 5-min of baseline and the last 5-min of drug application were used for quantification. Events were filtered online at 1 kHz and counted manually utilizing Minianalysis software (Synaptosoft, Leonia, NJ). At least 100 events per cell were acquired in 6 s blocks and detected using a threshold of 7 pA.

For the modulation of NMDAR currents, recordings were made at a +40 mV holding potential with DNQX (10 µM), the GABA_A_ antagonist picrotoxin (PTX, 50 µM) and TTX (1 µM) in the bath to isolate glutamate uncaging-evoked NMDAR currents and eliminate circuit effects evoked by glutamate uncaging. Glutamate uncaging was achieved by applying a single 150-ms pulse of UV light (356 nm) through the microscope objective every 20 s to a bath containing MNI-glutamate (50 µM, Tocris, Bioscience, Bristol, United Kingdom). NMDA components were measured as currents 20 ms after the peak. LED intensity was chosen to evoke responses at approximately half of maximal amplitude. Evoked NMDAR currents were recorded until their amplitudes were stable for at least 5 min, at which point PD-128907 (1 µM) or SKF-81297 (10 µM) was added to the bath for 10 min. SCH-39166 (1 µM) was already present in the bath to pharmacologically antagonize D1R signaling. Timecourse graphs were generated by averaging raw NMDAR measurements in 1-min bins and expressing each point as a percentage of the average of the 5-min baseline. The averaged baseline and last 5-min of drug application were used for quantification. Bath application of AP5 at the end of the experiment abolished glutamate uncaging-evoked currents, confirming that evoked currents were mediated by NMDARs. For validation of *Drd1a* KO, AAV8-Syn-GFP-Cre was injected into the NAc to genetically ablate *Drd1a* and CTB594 was delivered in the LH or VTA to visualize putative NAc D1R-expressing MSNs. Whole-cell recordings of NMDAR currents were performed on GFP-positive and CTB594-positive neurons as detailed above.

After recordings, images of the recording pipette were acquired for post hoc estimation of recording location within using the same camera as above and using a 4x Nikon objective. For all pharmacological experiments, one neuron per slice was analyzed. In some cases, the slice was transferred to 4% paraformaldehyde overnight for post-hoc imaging on a confocal microscope. The Clampfit suite v11.0.3.03 (Molecular Devices) was used for data display, acquisition and storage.

### Behavior

Behavioral experiments were conducted during the dark cycle (between 10 am and 6 pm) unless otherwise specified. Mice were allowed to recover from surgery for at least 3 weeks before behavioral testing was conducted. Animals were matched according to age, sex and date of birth and single-housed at least one day before the start of testing. To minimize the effect of stress on behavioral outcome, mice were acclimated to the red lit and sound-proofed testing room for at least 30-min before the start of each assay.

Animals undergoing running behavioral testing were subjected to the following assays in this order: wheel running, running disks choice test, open field test, sucrose preference test, social interaction, light-dark box, and novel object recognition test (Extended Data Fig. 1b). Separate cohorts of mice were used for operant conditioning experiments. For pathway-specific D3R cKO, mice underwent testing in running wheel, running disk choice behavior, and operant procedures.

#### Wheel running

Voluntary running was performed in mice with free access to a running wheel in their cage. At least 24-h before the start of the experiment, mice were singly housed in clean, standard cages to habituate them with social isolation in activity recording chambers. At the start of the light cycle (*i.e.*, 8 pm), mice were transferred into cages that contained a running wheel connected to an infrared sensor that recorded beam breaks on the wheel to calculate distance traveled (ACT-551-FIL-MS-SS, Coulbourn Instruments, Holliston, MA). Animals were provided with provided with ad libitum access to food and water, and running activity was monitored for 60 consecutive hours. Data was collected every 5-min from each mouse using Clocklab (Actimetrics, Wilmette, IL).

#### Running disks test

We designed an effort-related choice-behavior task to disentangle motivational running behavior. Mice had to choose between two disks, one that was fixed, where animals could not run on it, and another one that was freely moving, where animals experienced reward by running on it. Disk were angled at approximately 30° relative to the floor. Each session consisted of baseline and test phases. During baseline, the animal was placed in the empty open-field arena and allowed to explore the environment for 10 min. After baseline, the fixed and freely moving disks were inserted into the arena and disk-running activity and general locomotion were recorded for 3 hrs. There was no interruption between baseline and test phases. Visual (circles or stripes) and odor cues (ddH2O or 5% acetic acid) were attached adjacent to each disk to further facilitate the recognition of each area of the arena. Tracking data was analyzed offline using TopScan software (CleverSys, Inc., Reston, VA). The position of the mouse was defined as the zone in which both front paws and center were located.

#### Open-field test

Mice were placed singly in an open field arena (43.8 cm × 43.8 cm × 39.4 cm) to assess general locomotor activity and anxiety-like behavior. For the former, the total distance traveled was measured for 30 min. For the latter, the arena was divided into ‘center’ (23 x 23 cm) and ‘edges’ zones and the percentage of time spent in the center of the arena was measured. For both analyses, TopScan (CleversSys, Inc.) video tracking software was used to score the movement and location of the animals.

#### Sucrose preference test

Hedonic reward-seeking was measured using the sucrose preference test. Single-housed mice were first habituated by being placed a standard cage (45 × 27 × 15cm) that contained two bottles of tap water. Water intake was measured by weighing the bottles four hours after the start of the habituation period. The following day, one of the bottles was replaced with a 1% sucrose solution (wt/vol), and animals were again given a free choice between the two bottles. The total amount of tap water and sucrose consumed was recorded by again weighing the bottles after four hours. Sucrose preference was calculated as the amount of sucrose solution consumed relative to the total amount of liquid consumed and multiplied by 100 (sucrose solution intake/total intake) *100). To control for side preferences the location of the sucrose and water bottles was counterbalanced between cages. Sucrose preference testing occurred approximately 2 hrs after the start of the animal’s active cycle.

#### Light-dark box

The light-dark box test was performed to assess anxiety-like behavior. Mice were place in an open-field arena (43.8 cm × 43.8 cm × 39.4 cm) divided into two compartments connected by a small circular aperture (4 W x 5 H cm). One side was exposed to light in the room and the other was enclosed and dark. During testing, mice were placed in the lit side of the box, facing the wall farthest from the entrance to the dark side. Animals were allowed to explore the two compartments for 5 min. Videos were recorded with a camera positioned above the chamber. The latency to enter the dark compartment and the time spent in each side were quantified offline with TopScan software. Room lighting was measured with the aid of a lux meter during testing (∼100 Lux).

#### Novel object recognition test

The novel object recognition task assessed novelty-seeking behavior, capitalizing on rodents’ natural inclination to spend more time with a novel object than over a familiar one^108^. Object recognition testing was carried out in a plexiglass open-field box (43.8 cm × 43.8 cm × 39.4 cm) and comprised in two sessions: habituation and recognition. During the habituation session, animals were allowed to freely explore the environment, which contained two identical Lego® constructs in opposite corners of the box, for 10 minutes. Items were placed on a metal base to ensure they could not be moved or knocked over by the animals. Subsequently, mice were taken back to their home cages for an ITI of 1 hour and were reintroduced in the apparatus for the recognition test lasting ten minutes. In this 10-min session, one of the two objects used in the habituation phase was replaced with a novel object that was differently colored and shaped compared to the original, familiar object. The identity and position of the novel and familiar objects were counterbalanced across groups. Objects were thoroughly cleaned with water between phases to remove odor traces. Total spent time exploring each of the objects during both phases was quantified offline using TopScan software. The discrimination ratio was calculated as the time spent interacting with the novel object area divided by the time spent in a novel object area plus the time spent in the familiar object area.

#### Social interaction assay

For social interaction experiments, mice were temporarily moved to a target-free holding arena (56 × 24 x 24 cm) that contained two empty mesh pencil cups (5× 6.5 × 8 cm) in opposite corners of the arena. Animals were first allowed to freely explore the chamber for 2.5 min before a novel juvenile mouse (3-6 weeks) of the same sex and strain (to avoid mutual aggression) was placed into one of the holders. The test mouse was allowed to freely interact for 7.5 min, and video was recorded with a camera suspended above the arena. The ‘interaction zone’ encompassed a circular area projecting 6.5 cm around the pencil cup. The ‘corner zones’ encompassed a 9 cm × 9 cm area projecting from both corner joints opposing the wire-mesh enclosure. Social interaction was automatically scored with AnyMaze software (Stoelting, Kiel, WI) and defined as the ratio of time spent in the interaction zone with a juvenile mouse (*i.e.* in active contact with the intruder’s snout, flank, or anogenital area, grooming, 6.5 cm from the enclosure of the pencil cup) over time spent with the target absent.

#### Operant conditioning

Prior to initiating operant procedures, mice were weighed to the nearest 0.1 g to determine a baseline for free-feeding body weight. Mice were singly housed and maintained under food restriction to achieve 85-90% of their free-feeding body weights for 3 days before and throughout the experiments, which motivated them to perform the behavioral task. Mice were weighed daily and fed 1-hour after their daily behavioral sessions with 2 g of standard laboratory chow. Animals had free access to water throughout. Operant procedures were conducted 7 days per week over a 4-wk period.

Chocolate pellet self-administration was used to examine reward-related behaviors and took place in sound-attenuated mouse operant chambers (ENV307A-CT, Med-Associates, Fairfax, VT, PC 5 software). Chambers were equipped with two retractable levers and a reward pellet dispenser. One lever was designated as ‘active’ and was paired with the delivery of chocolate pellets (20 mg, Bio-Serv, Flemington, NJ), while the other lever was designated ‘inactive’ awhere lever-pressing had no consequence on reward delivery. The assignment of active and inactive levers was counter-balanced between mice. There was a 5-s time-out after every pellet delivery during which lever pressing did not trigger any delivery. The location of active and inactive levers was counterbalanced across animals and chambers were kept dark (house light off) during all sessions. A house light was positioned above the levers and a fan was present to maintain ventilation throughout testing. Before the start of the conditioning experiments, animals first underwent two pretraining sessions in which chocolate pellets were delivered at a random interval schedule (mean of 45 s, range 4–132 s), where pressing in either lever had no consequence on reward delivery. Days 1-6 of operant training were conducted on a fixed-ratio 1 (FR1) schedule, and days 7-12 were conducted on an FR5 schedule, where mice had to press the active lever 1 and 5 times, respectively, to earn a reward. Mice underwent testing in one session each day. All FR sessions lasted 45 min.

##### Operant Choice task

Our operant choice task was modified from previous reports^46,47^. After 6 days of testing on FR5 schedule, mice underwent testing on the operant choice task. First, mice experienced a session where they were presented with a food receptacle containing freely-available regular laboratory chow or chocolate pellets in the corner of the operant chamber opposite of the wall containing the active lever and operant reward receptacle. This was done to expose the animals to freely-available food in the operant chambers. During this session, active and inactive levers were retracted. Freely-available reward exposure sessions with regular laboratory chow or chocolate pellets were conducted on different days prior to choice testing. Mice were subsequently tested in a choice task, where they could either lever-press on an FR5 schedule to obtain a highly palatable food (20 mg chocolate food pellets) or consume the freely available food over the grid located on the opposite side of the operant chamber in a food receptacle. Choice sessions with regular lab chow or chocolate pellets were conducted on different days. After a choice session, a retrain session where only chocolate pellets delivered on the FR5 schedule were available was presented to the mice. The number of lever-presses, quantity of freely-available food (standard chow or chocolate pellet) consumed, total amount of food consumed (pellets plus chow) and the amount of chocolate pellets obtained by lever pressing were recorded.

##### Progressive ratio operant tasks

After FR schedule sessions, a progressive ratio (PR) schedule of reinforcement for chocolate pellets was used to assess the motivation to work for chocolate pellets. To familiarize animals with a schedule requiring more effort, a PR3 schedule was used for 3 days. Under this schedule, response increments linearly increase by three lever presses (3, 6, 9, 12, 15, etc.) for delivery of each subsequent food pellet. Animals subsequently underwent 4 days of training on a PR7 schedule to assess motivation under a schedule with higher demands. During PR testing, sessions continued until 5 min had elapsed without the animal responding in the active lever. In each PR session the break point (the final number of responses an animal completes where a reward is delivered), was recorded. All PR sessions ended after 3 hours or until 5 mins elapsed without a response in the active lever.

#### Platform-mediated avoidance task

This modified tone-shock conditioning experiment tested for reinforcement of avoidance behavior. Experiments were conducted in sound-attenuated fear conditioning chambers (30 cm length × 25 cm width × 25 cm height, Med Associates) that were illuminated with red light. The floor of the chamber was composed of a metal grid that delivered an electric foot shock. All tests began with a 2-min baseline habituation. Following the baseline period, 20 pairings of a conditioned stimulus (CS, 30 sec, 80-dB, 4 kHz noise) co-terminating with an unconditioned footshock stimulus (US, 0.4 mA, 2 sec). The ITI between trials was 40-60 sec. Animals could avoid the shock by jumping onto a square Plexiglas platform (8 x 8 x 0.33 cm) located on a corner of the chamber that was fixed to the shock floors. Each experiment lasted for two days with one 30-min session conducted each day. Day 2 consisted of identical stimuli presentations as day one. Between experiments, shock grids and floor trays were cleaned with soap and water, and chamber walls were cleaned with wet paper towels. Tracking data was acquired and analyzed using AnyMaze software (Wood Dale, IL). This task was modified from previous reports to adapt it to mice^77,78^.

### D3R pharmacological inactivation

Microinjection of the D3R antagonist SB-277011A (Tocris) was used to pharmacologically inhibit D3R signaling in the NAc. Animals were allowed to recover for 4 weeks after surgery before habituation to the microinjection procedure. For 2 days prior the start of the experiments, mice were habituated to handling and cannula manipulation. On the experimental day, 350 nL of 10 µM SB-277011A (1.79 ng, dissolved in 1% DMSO) or vehicle (1% DMSO) were bilaterally infused into the NAc. This was accomplished using 33-gauge injector cannulas connected to a syringe pump (UMP3, World Precision Instruments) with PE20 tubing that protruded 1 mm beyond the tip of the 26-gauge guide cannula All microinjections were delivered over the course of 1 min. After infusion, injectors were left in place for 2 min to allow for complete drug diffusion. For D3R functional disconnection procedures, mice were habituated 2 days before the experiment by receiving an infusion of vehicle (1% DMSO). 5 min after the end of the intra-NAc injection, animals were placed in the open-field arenas and running disk choice testing was conducted. Cannula placements were verified by histology after injection of 300 nL per hemisphere of red fluorescent Retrobeads (Lumafluor).

### D1R-NMDAR functional disconnection

For 2 days prior the start of D1R-NMDAR procedures, mice were habituated to handling and cannula manipulation. On the experiment day, 350 nL per hemisphere of the competitive NMDAR antagonist AP5 (700 ng per infusion, dissolved in 0.9% saline) were infused according to the same procedure as D3R pharmacological inactivation. Cannula placements and spread of infused drug were verified by histology after injection of 300 nL per hemisphere of red fluorescent Retrobeads (Lumafluor).

### Chemogenetic inhibition of D3R-expressing NAc MSNs

*Drd3*-Cre mice undergoing inhibitory DREADD testing were allowed to recover for 4weeks following viral injection before the start of behavioral testing. On the day prior to testing, mice received an intraperitoneal injection of saline to habituate them with the injection procedure. On the day of the experiment, the DREADD agonist clozapine N-oxide hydrochloride (CNO; Enzo Life Sciences, East Farmingdale, NY) was administered intraperitoneally 30 min before the running disks test. CNO solutions were intraperitoneally injected at 0.1 ml solution per 10 g of mouse for a final concentration of 1 mg/kg (in sterile saline).

### Drugs

(+)-PD 128907 hydrochloride (PD128907, Tocris), ML417 (Sibley Lab, NINDS) and SKF-81297 (Tocris) were dissolved in distilled water. SCH-39166 (Tocris) was dissolved in DMSO. SB-277011A (Tocris Biosciences) was dissolved in 1 % DMSO and administered at 10 µM. AP5 (Abcam, Cambridge, UK) was diluted in sterile 0.9 % saline. For electrophysiology experiments, 1 mM PD128907 and 1mM SCH-39166 were diluted to a final concentration of 1 µM in aCSF and 10 mM SKF-81297 was diluted to a final concentration of 10 µM in aCSF.

### Quantification and Statistical Analysis

Data analysis was performed using GraphPad Prism 9, (GraphPad Software, Inc., La Jolla, CA). Paired t-tests were used for within-group comparison of two treatments and an unpaired test was used for comparison between two groups. Differences across more than two groups were analyzed with either a one-way analysis of variance (ANOVA) with Tukey’s or Dunnet’s multiple comparison post-hoc test, a two-way ANOVA for data with Tukey’s or Sidak’s multiple comparison post-hoc test for two independent variables, or a two-way repeated-measures (RM) ANOVA for data with two independent variables and multiple measurements from the same subject. ANOVAs were followed by post hoc tests with multiple comparisons correction. In the case of datasets with missing values, we analyzed the data instead by fitting a (one-way) mixed model as implemented in GraphPad Prism 9.0. Kolmogorov–Smirnov test was used for cumulative probability plots. P values for linear regressions were calculated by using Pearson’s correlation. For each experiment, the values and definitions of sample size (n) are explicitly explained in Extended Data Table 1. Statistical significance was defined as *P < 0.05, **P < 0.01, ***P < 0.001, ****P < 0.0001. R^2^ represents a Pearson’s correlation coefficient. Results are shown as mean ± SEM unless stated otherwise. Error bars represent SEM. See Table S1 for detailed statistics.

### Data and code availability

Data and analysis code reported in this paper is available from the lead contact upon reasonable request.

## Supporting information

Supplemental Table 1 Statistics

Supplemental Table 2 Key Resources

## Acknowledgements

This work was supported by the National Institute of Mental Health Intramural Research Program (ZIA MH002970-04), a Brain and Behavior Research Foundation NARSAD Young Investigator Award to HAT, NIH Center for Compulsive Behavior Fellowships to RFG and HEY, Department of Defense PRMRP Investigator-Initiated Award (PR141292) to Z.F., Department of Defense PRMRP Expansion Award (PR210207) to Z.F., John F. and Nancy A. Emmerling Fund of The Pittsburgh Foundation (Z.F.), a NIH Post-Doctoral Research Associate Training Fellowship to RFG, and a Medical Research Scholars Program Fellowship to CTL. The authors would like to thank Sarah Williams and Jonathan Kuo of the NIMH Systems Neuroscience Imaging Resource and Dr. Samer Hattar for their microscopy support. The authors also acknowledge Dr. Yogita Chudasama and members of the NIMH Rodent Behavior Core for behavioral equipment support and Dr. Samer Hattar for sharing running wheels. The authors also thank Drs. Claudia Schmauss, Jonathan Javitch, and Marcelo Rubinstein for the design of the *Drd3*^fl/fl^ mouse strain as well as Dr. Siu-Pok Yee and the Center for Mouse Genome Modification at UConn Health for the construction of the *Drd3*^fl/fl^ mice. We additionally acknowledge Dr. Veronica Alvarez and members of the Alvarez laboratory including Drs. Lauren Dobbs and Miriam Bocarsly for all of their invaluable help in making the *Drd3*^fl/fl^ mice congenic with the C57BL/6J genetic background. We would like to thank members of the Unit on Neuromodulation and Synaptic Integration and Drs. Mario Penzo, Marco Pignatelli, Federica Lucantonio, Eastman Lewis, Detlef Vullhorst, and Andres Buonanno for discussions and critical reading of the manuscript.

## Author information

Conceptualization, J.E.T. and H.A.T.; Methodology, J.E.T., T.B.U., and H.A.T, Software, J.E.T., R.J.F. and S.R.; Validation, J.E.T.; Formal Analysis, J.E.T. and V.T.; Investigation, J.E.T., H.E.Y., H.A.T., Z.F., T.W., C.T.L., M.A., H.W. and R.J.F.; Resources, Z.F., A.E.M., D.R.S. and H.A.T.; Data Curation, J.E.T.; Writing – Original Draft, J.E.T.; Writing, Revision and Editing, J.E.T., Z.F. and H.A.T.; Visualization, J.E.T.; Supervision, R.M. and H.A.T.; Project Administration, J.E.T. and H.A.T.; Funding Acquisition, Z.F. and H.A.T.

---

**Extended Data Figure 1.**
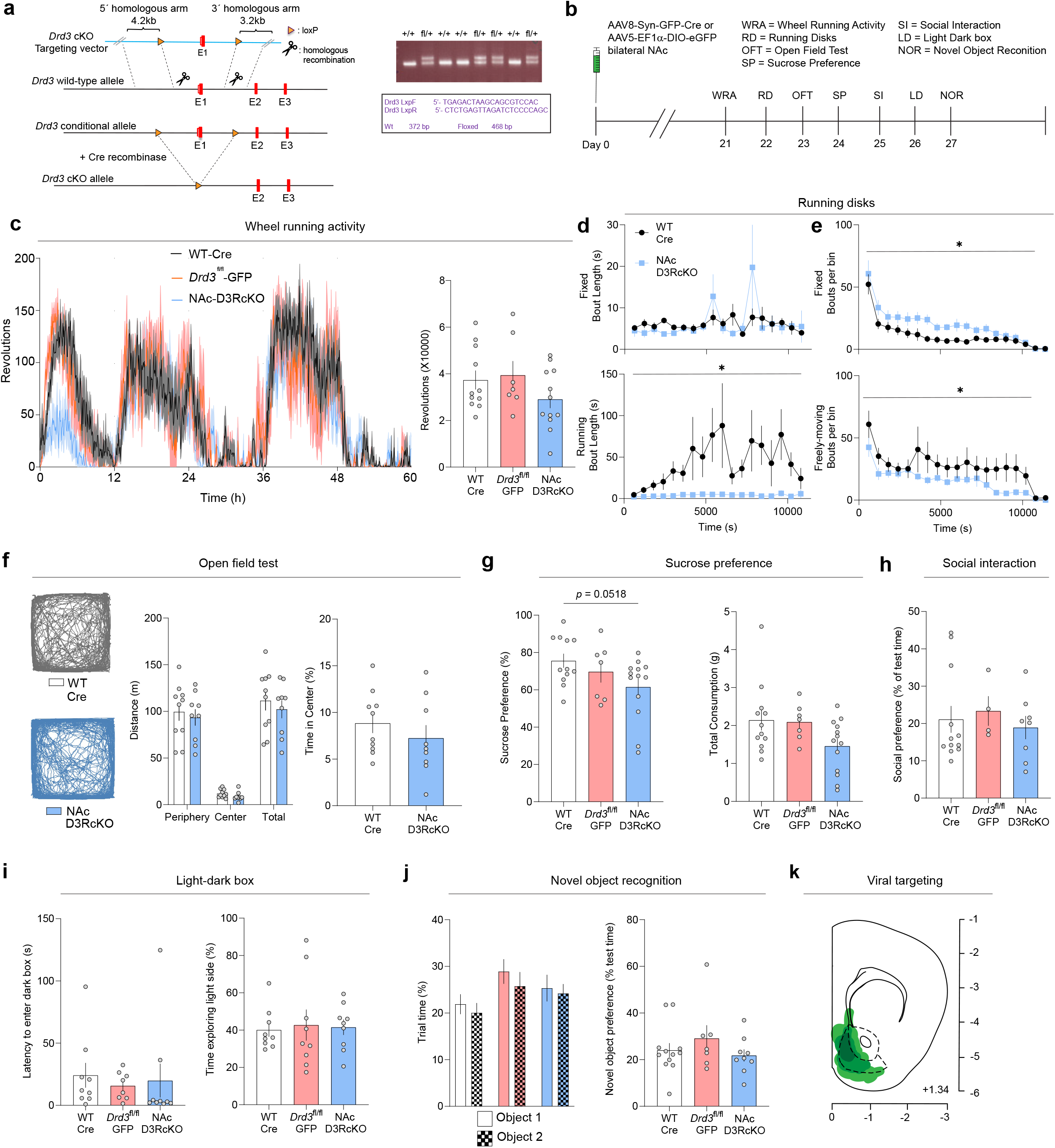
NAc D3Rs does not affect locomotion, anhedonia, social reward, anxiety or novel object-recognition, Related to Figure 1. a. (Left) Schematic depicting the strategy used to generate the *Drd3*^fl/fl^ conditional-knockout (cKO) mouse strain. LoxP cassettes flanking the exon 1 were inserted in the coding region of the *Drd3* gene that encodes the D3R protein using homologous recombination approaches. (Right) Confirmation of the inserted LoxP sites within the chimeric animals by using PCR strategies (see primers in purple). The successful insertion of the LoxP elements was confirmed by the presence of two PCR bands (fl/+ lanes) versus a single band in the wildtype littermates (+/+ lanes). b. Experimental timeline of behavioral experiments in mice involving running behavior. c. (Left) Timecourse of wheel-running activity for the entire duration of the experiment (60 hrs) (Right) Quantification of total revolutions per group during the entire 60 hr period. d. Length of visit (bout) to the fixed disk (top) or freely-moving disk (bottom, putative running bout) across the running disks session. e. Number of entries into the fixed (top) or freely-moving disk (bottom) across the running disks session. f. (Left) Representative cumulative locomotion traces of open field activity in WT-Cre and NAc-D3RcKO groups. (Middle) Quantification of cumulative distance traveled. (Right) Percentage of time spent in the center during the open field test. g. (Left) Percentage of sucrose preference. (Right) Overall intake in the sucrose preference test. h. Social preference as reflected by the % time (test-habituation) spent interacting with a novel, juvenile mouse. i. Anxiety-like behavior as represented by the latency to enter dark chamber in the light–dark box (left) or time spent in the light side of the light-dark box (right). j. (Left) Time spent interacting with each of the objects during the baseline period. (Right) Preference for the novel object over a familiar one during the discrimination test. k. Schematic of combined viral spread map of local *Drd3* cKO. Dark green indicates animal with most restricted expression and lighter green indicates animals with broader pattern of viral spread.

**Extended Data Figure 2.**
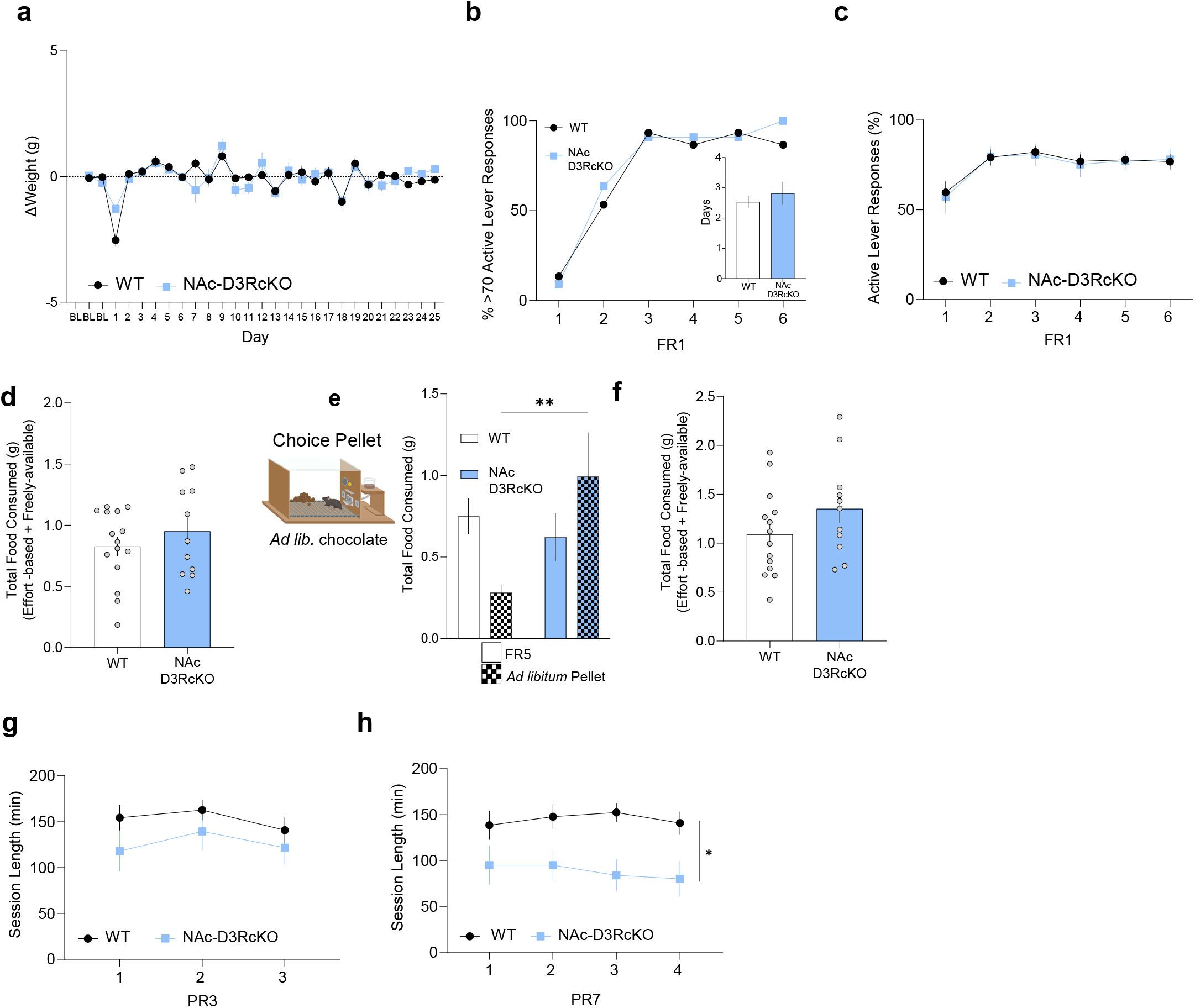
NAc D3R regulates motivation towards working for rewards, but does not affect weight, or FR acquisition schedules of reinforcement, Related to Figure 1. a. Body weight changes at baseline before food deprivation or across the overall duration of operant conditioning procedures in Figures 1 and S1. b. FR1 acquisition, as measured by the percentage of animals reaching the criteria of 70 active lever responses per session. (Inset) Days required to acquire FR1 (criteria for acquisition was 70 responses in a 45 min session). c. Percentage of active lever responses during FR1 schedules. d. Total food consumed in choice chow session. e. (Left) Diagram of the FR5 choice pellet session. (Right) Amount of food consumed, which is represented as FR5 effort-based or freely-available chocolate pellets. f. Total food consumed in FR5 choice pellet session. g. PR3 session length. h. PR7 session length.

**Extended Data Figure 3.**
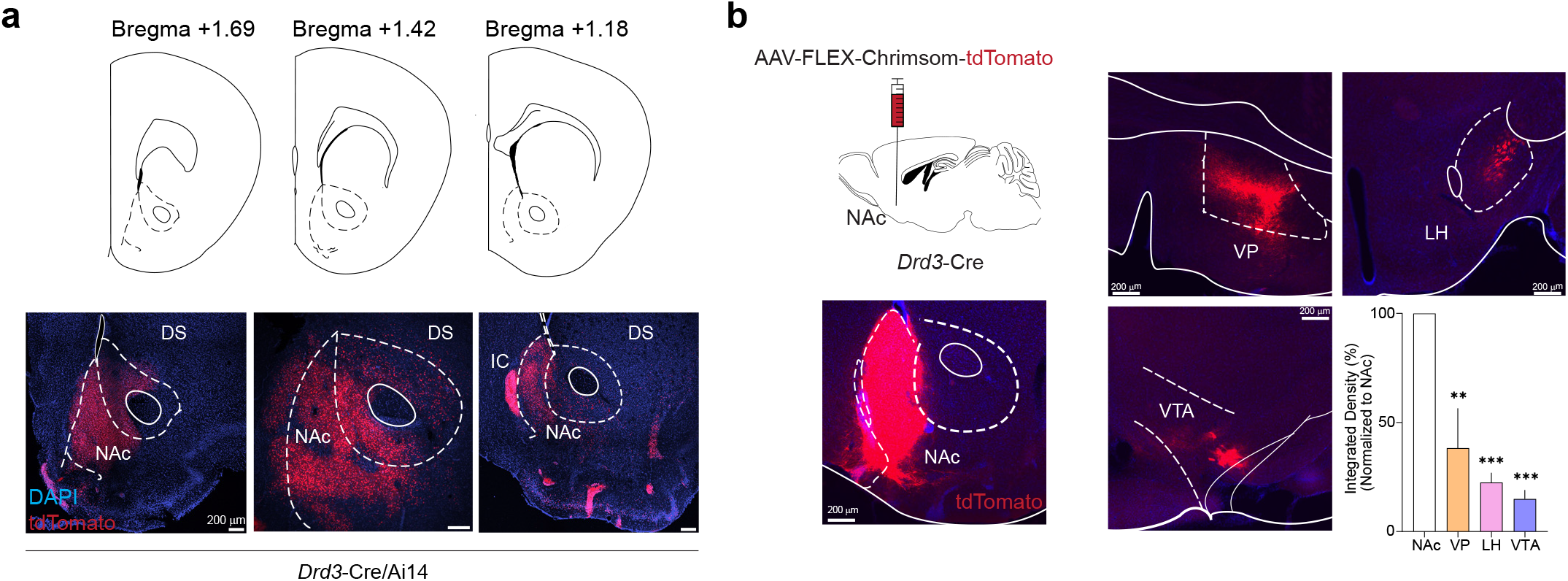
D*rd3*-Cre lines provide genetic access to D*rd3* expression in the NAc and outputs of D3-MSNs, Related to Figure 2. a. *Drd3* expression in the NAc of *Drd3*-Cre/Ai14 mice. Coronal diagrams depicting the region analyzed and confocal images showing the *Drd3* expression pattern in the NAc along the dorsal-ventral and rostral-caudal axis (bottom). DS = dorsal striatum; IC = islands of Calleja b. (Left) A virus encoding the anterograde tracer (AAV1-Syn-FLEX-Chrimsom-tdTomato) was targeted to the NAc in *Drd3*-Cre mice that express tdTomato in a Cre-dependent manner. (Right) Representative images and quantification of tdTomato^+^ fibers in NAc, VP, LH and VTA regions.

**Extended Data Figure 4.**
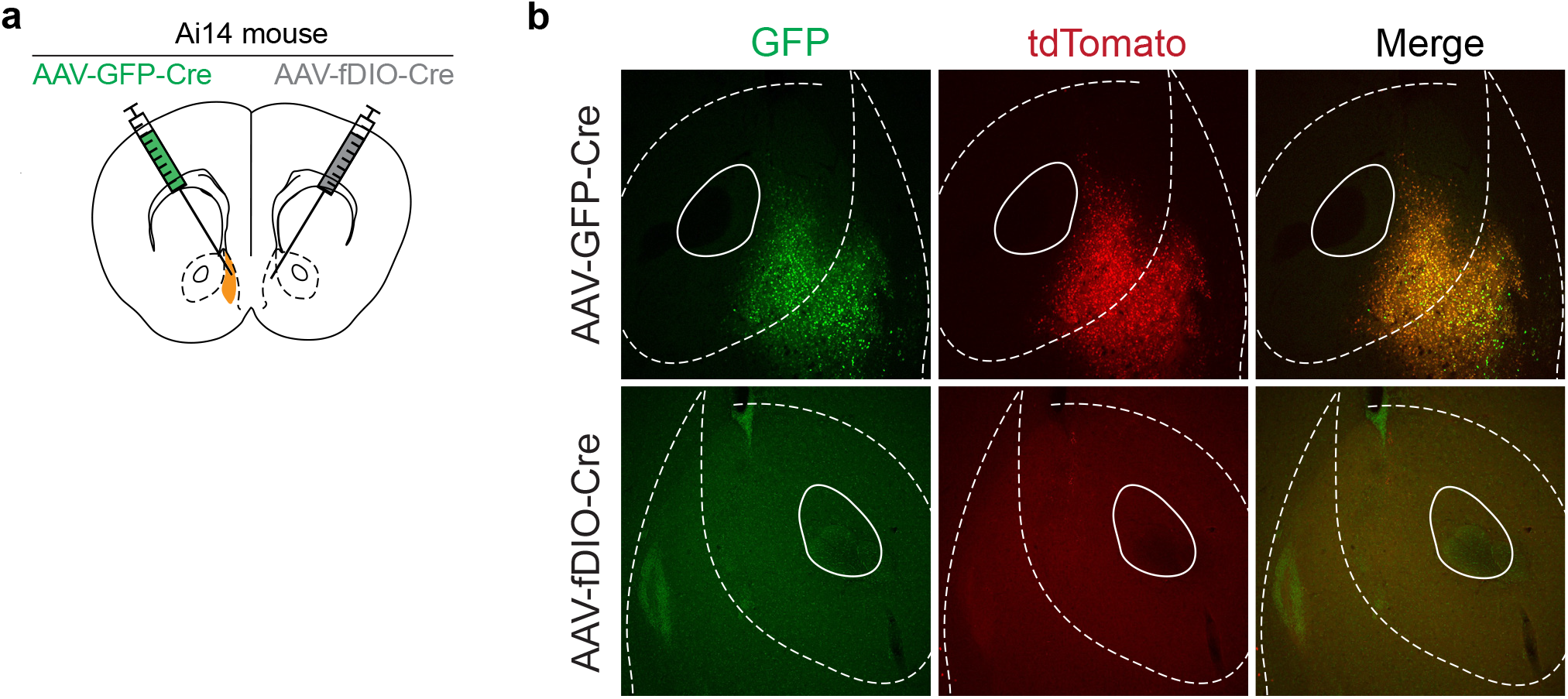
Validation of lack of Cre leakage in our intersectional approach for pathway-specific NAc D*rd3* cKO, Related to Figure 3. a. Scheme of the experimental design to validate the pathway-specific strategy for the pathway-specific cKO of NAc *Drd3*. AAV8-Syn-GFP-Cre and AAV9-EF1a-fDIO-Cre were injected in contralateral hemispheres of Ai14-tdTomato reporter mouse. b. Representative images showing colabeling of GFP and tdTomato in hemisphere injected with AAV-GFP-Cre (top) and lack of tdTomato expression in hemisphere injected with AAV-fDIO-Cre (bottom).

**Extended Data Figure 5.**
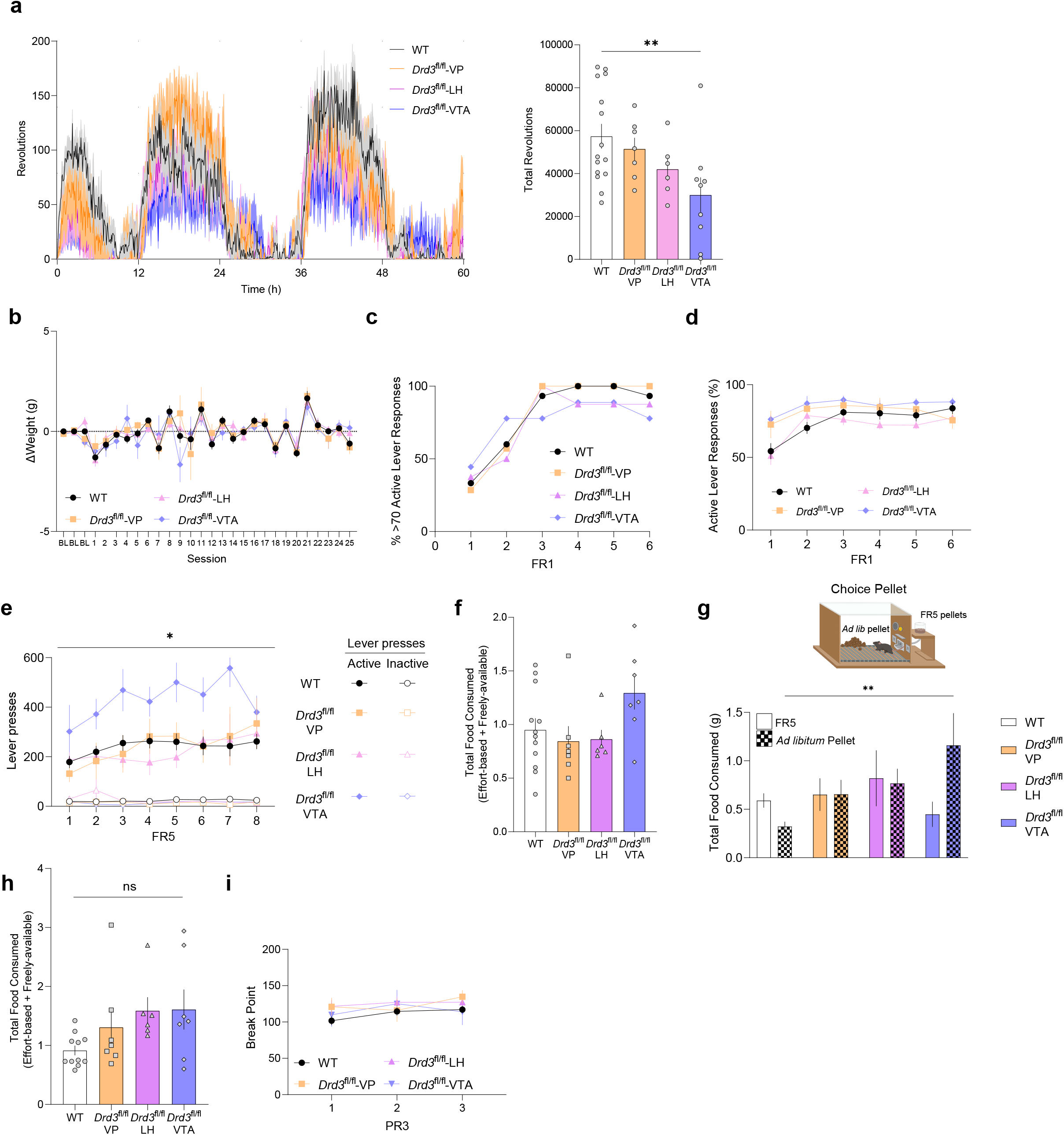
NAc D3Rs provide value to running and working for reward, but does not regulate acquisition of reinforcement, Related to Figure 3. a. Wheel running activity for pathway-specific deletion of NAc D3Rs. (Left) Timecourse of running activity during the complete 60-hr experiment. (Right) Quantification of total revolutions during the 60-hr period. b. Body weight changes at baseline before food deprivation (*p*= 0.86) or across the overall duration of operant conditioning procedures in Figure 3. c. FR1 acquisition, as measured by the percentage of animals reaching the criteria of 70 active lever responses per session. d. Percentage of active lever responses during FR1 schedules e. Lever presses during FR5 acquisition sessions. f. Total food consumed in choice chow session. g. (Top) Diagram of the FR5 choice pellet session. (Bottom) Amount of food consumed which is represented as either FR5 effort-based (solid) or freely-available chocolate pellets (checkered). h. Effects of region-specific D3R cKO on total consumption of chocolate pellets in a FR5 choice pellet session. i. Break points during the PR3 reinforcement schedule for VP-, LH-, and VTA-specific D3R cKO versus WT controls.

**Extended Data Figure 6.**
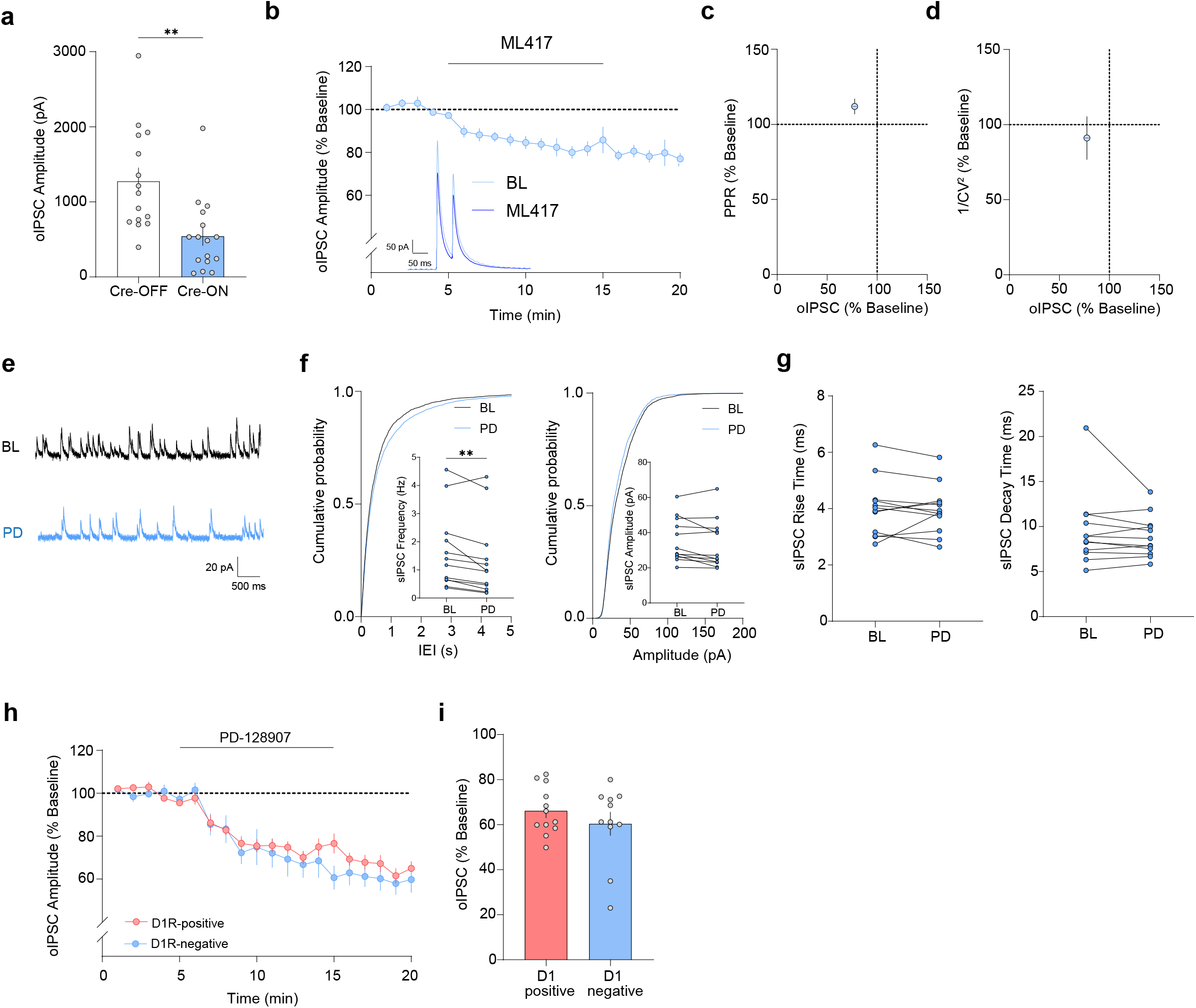
NAc D3R decreases GABA release probability presynaptically onto both D1- and D2-MSNs, Related to Figure 4. a. Mean baseline oIPSC amplitude (pA) for Cre-OFF (white bar) and Cre-ON (blue bar) evoked in NAc MSNs. b. Timecourse of oIPSCs in NAc MSNs before, during and after bath application of the D3R-selective agonist ML417 (1 µM) in *Drd3*-Cre mice expressing Cre-dependent ChR2 (blue). (Inset) Representative oIPSC traces recorded in NAc MSNs before and after bath application of ML417. c. Paired-pulse ratio (PPR, % baseline) versus oIPSC (% baseline) after ML417 application. d. Coefficient of variation (1/CV^2^, % baseline) versus oIPSC (% baseline) after ML417 application. e. Representative traces of sIPSCs during baseline (BL, black) and after bath-application of the D3R-selective agonist PD-128907 (PD, blue). f. Cumulative probability of sIPSC inter-event interval (left) and amplitude (right) recorded from NAc MSNs. (Inset) Quantification of frequency and amplitude of sIPSCs events during baseline and after PD-128907 application g. Rise (left panel) and decay time (right panel) of sIPSC events during baseline and after PD-128907 application h. Time-course of oIPSCs before, during and after bath application of PD-128907 in NAc D1R-positive and D1R-negative NAc neurons (putative D2-MSNs) in the Cre-ON condition. (Inset) Representative oIPSCs traces recorded in NAc D1- and D2-MSNs before and after bath application of PD-128907. i. Bar-graph quantification of oIPSC inhibition after PD-128907 application in D1R-positive and D1-negative NAc MSNs.

**Extended Data Figure 7:**
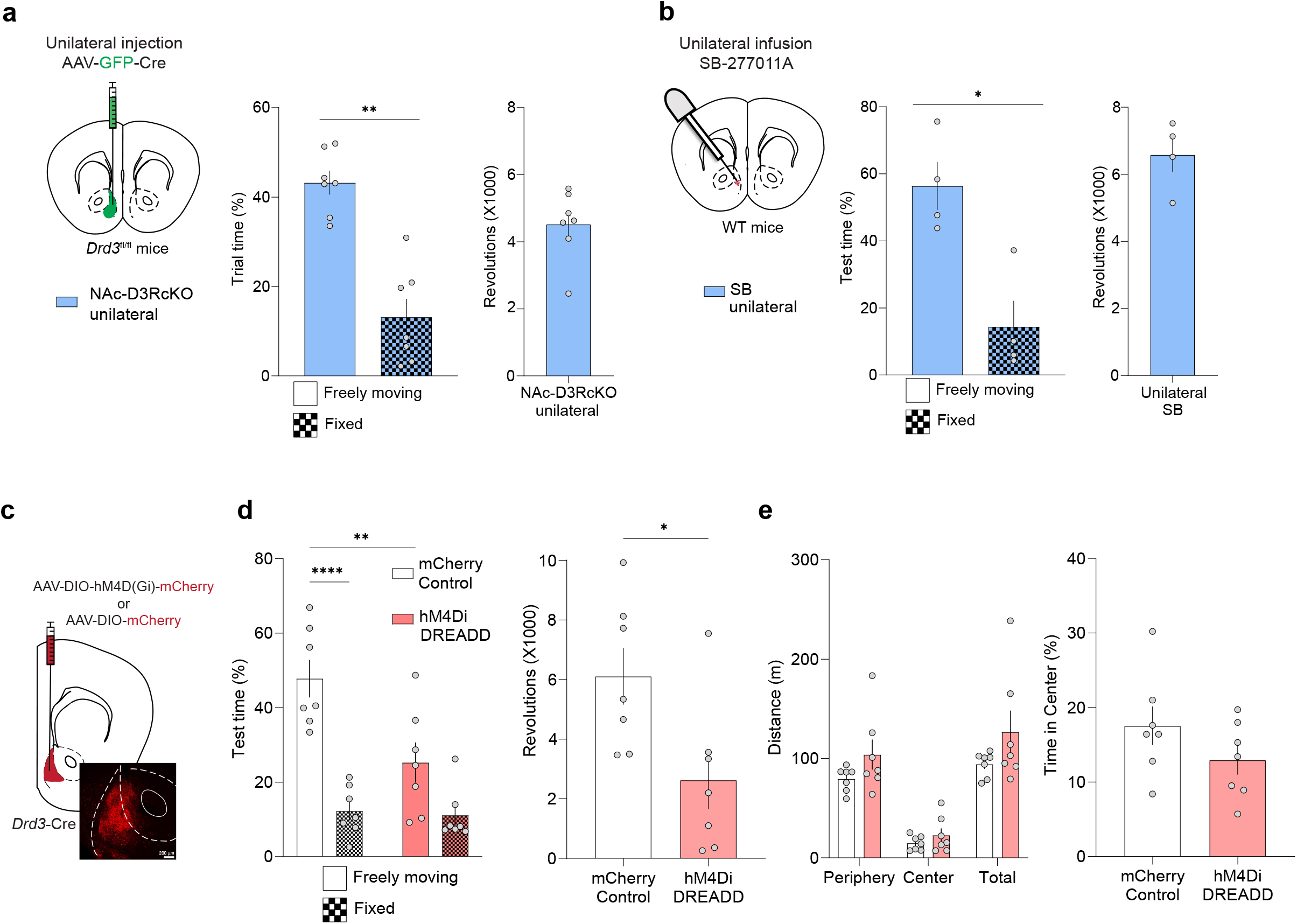
Unilateral suppression of D3R signaling does not disrupt motivated running behavior, Related to Figure 5. a. Quantification of time spent in the freely-moving and fixed disk and revolutions for *Drd3*^fl/fl^ mice expressing unilateral Cre recombinase. b. Same as in (a) but for WT control mice unilaterally infused with SB-277011A into the NAc. c. Schematic (top) and representative image (bottom) of viral expression of AAV5-hSyn-DIO-mCherry in the NAc of *Drd3*-Cre mice d. (Left) Percentage of time spent on fixed or freely-moving disk during the running disk task for mCherry and HM4Di groups. (Right) Running behavior as represented by the number of revolutions registered in the freely moving disk. e. (Left) Quantification of cumulative distance traveled in each zone during the open-field test. (Right) Percentage of time spent in center.

**Extended Data Figure 8:**
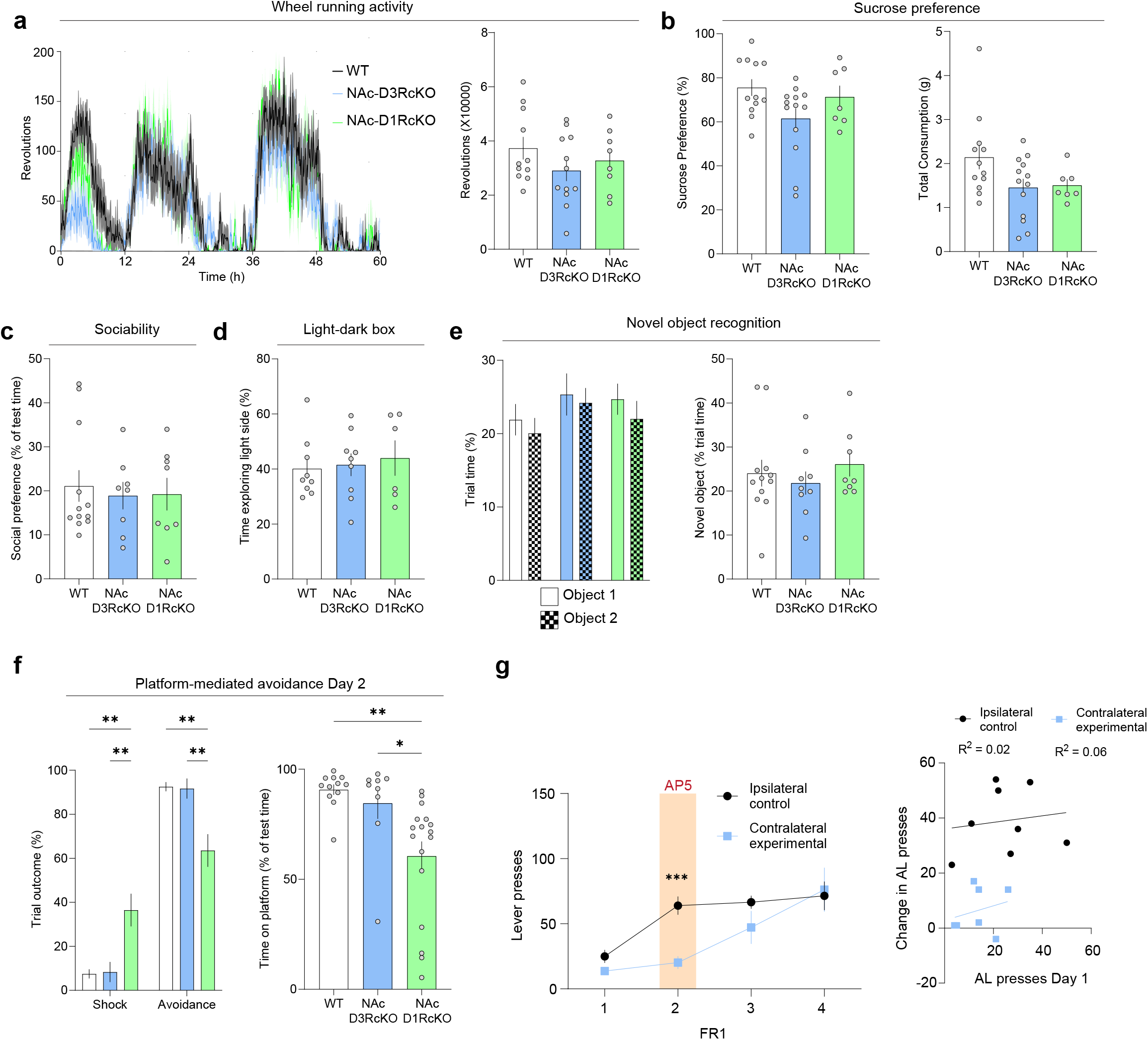
NAc D1Rs do not mediate motivated, anxiety-like or social reward or sucrose preference, Related to Figure 6 and Figure 7. a. (Left) Time course of wheel-running activity across the entire duration of the experiment (60-hrs) for WT, NAc-D3RcKO and NAc-D1RcKO groups in 5 min bins. (Right) Quantification of total revolutions across the 60 hr period.. b. (Left) Percentage of sucrose preference for WT, NAc-D3RcKO and NAc-D1RcKO groups. (Right) Overall water and sucrose intake. c. Social preference as reflected by the % time (test-habituation) spent interacting with a novel, juvenile mouse. d. Anxiety-like behavior as represented by the time spent in the light side of the box. e. Preference for the novel object over a familiar one during the discrimination test. f. (Left) Quantification of trial outcome (avoidance or shock responses) upon re-exposure to the platform-mediated avoidance task on day 2. (Right) Overall time spent on platform (Day 2) as percentage of test time. g. (Left) Absolute number of active lever responses in FR1 sessions for D1R-NMDAR disconnection experiments. AP5 infusion was performed on Day 2 of FR1. (Right) Correlation between number of AL presses on Day 1 and change in active lever presses on Day 2 (AP5 challenge).

