## Supplemental Table 2 Key Resources for "Dissociable control of motivation and reinforcement by distinct ventral striatal dopamine receptors"

**Extended Data Table 2. Key Resources**

| **REAGENT or RESOURCE** | | | **SOURCE** | **IDENTIFIER** |
| --- | --- | --- | --- | --- |
| Virus Strains | | | | |
| rAAV1-hSyn1-FLEX-ChrimsomR-tdTomato | | | UNC Viral Vector Core | Lot# AV6554B |
| rAAV2/9-phSyn1(S)-FLEX-tdTomato-T2A-Synaptophysin-eGFP-WPRE | | | Boston Children’s Viral Vector Core | Custom |
| rAAV5-EF1α-DIO-hChR2(H134R)-eYFP-WPRE | | | UNC Viral Vector Core | Lot# AV4313-2A |
| rAAV1-EF1α-DO-hChR2(H134R)-eYFP | | | NIDA Genetic Engineering and Viral Vector Core | Lot# AAV897 |
| rAAV1-CAG-FLEX-tdTomato | | | UNC Viral Vector Core | Lot# AV5328B |
| rAAV8-hSyn1-GFP-Cre | | | UNC Viral Vector Core | Lot# AV5053D |
| rAAV1-EF1α-eGFP | | | Addgene | 100043 |
| rAAV9- EF1α-fDIO-Cre | | | Addgene | Cat No. 121675 |
| CAV-Flp-GFP | | | Institut de Génétique Moléculaire de Montpellier CNRS UMR | N/A |
| rAAV10-Syn-DIO-mCherry | | | UNC Viral Vector Core | N/A |
| rAAV10-hSyn-DIO-HM4D(Gi)-mCherry | | | Addgene | 112677 |
| rAAV- EF1α-DIO-eGFP | | | GEVVC @ NIDA | AAV-2015-06-23-B |
| Chemicals, Peptides, and Recombinant Proteins | | | | |
| DNQX disodium salt | | | Abcam | Cat No. Ab120169 |
| D-AP5 | | | Tocris | Cat. No. 0106 |
| Picrotoxin | | | Tocris/Abcam | Ab120315(Abcam) |
| MNI-caged Glutamate | | | Tocris | Cat. No. 1490 |
| SCH-39166 hydrobromide | | | Tocris | Cat. No. 2299 |
| SKF 81297 hydrobromide | | | Tocris | Cat. No. 1447 |
| (+)-PD 128907 hydrochloride | | | Tocris | Cat. No 1243 |
| ML417 | | | Sibley Lab | N/A |
| SB-277011A dihydrochloride | | | Tocris | Cat. No. 4207 |
| Dimethyl Sulfoxide (DMSO) | | | Sigma-Aldrich | Cat. No. D8418 |
| Dustless precision pellets, chocolate | | | Bio-serv | F05301 |
| Red and green retrobeads IX | | | Lumafluor | R170 |
| DAPI Fluoromount-G | | | SouthernBiotech | Cat No. 0100-20 |
| Paraformaldehyde | | | Electron Microscopy Sciences | N/A |
| Clozapine-N-Oxide | | | Enzo Biosciences | BML-NS105-0025 |
| CTB-594 | | | Invitrogen | Cat. No. C34777 |
| Critical Commercial Assays | | | | |
| RNAscope Fluorescent Multiplex Reagent Kit 2.0 | | | ACDBio | Cat No. 320850 |
| RNAscope Probe against Mm-*Drd3* | | | ACDBio | Cat No. 447721 |
| RNAscope Probe against Mm-*Drd1a* | | | ACDBio | Cat No. 404691 |
| RNAscope Probe against Mm-*Drd2* | | | ACDBio | Cat No. 406501 |
| RNAcope Probe against *Cre recombinase* | | | ACDBio | Cat No. 474001 |
| Experimental Models: Organisms/Strains | | | | |
| Mouse: *Drd3*-Cre: B6.FVB(Cg)-  Tg(*Drd3*-cre)KI196Gsat/Mmucd | | | GENSAT/MMRC | Stock: Tg(Drd3-cre)KI196Gsat/Mmucd |
| Mouse: *Drd3*-Cre/Ai14 TdTomato reporter | | | Gift from Dr. Charles Gerfen |  |
| Mouse: *Drd3*^fl/fl^ | | | Gift from Dr. Zachary Freiberg |  |
| Mouse: *Drd1a*^fl/fl^ | | | Jackson Laboratories | Cat. No. 025700 |
| Mouse: *Drd1a*-tdTomato | | | Jackson Laboratories | Cat No. 016204 |
| Mouse: *Drd1a*-tdTomato/*Drd3*-Cre | | | NIH | N/A |
| Mouse: C57BL/6J | | | Jackson Laboratories | JAX#000664 |
| Software and Algorithms | | | | |
| Cell Profiler | | | Cell Profiler  (Broad Institute; Harvard and MIT) | <https://cellprofiler.org/> |
| ImageJ (FIJI) | | | NIH | <https://fiji.sc/> |
| GraphPad Prism 9 | | | GraphPad Software | <https://www.graphpad.com/> |
| Synapse | | | Tucker-Davis Technologies | <https://www.tdt.com/files/manuals/SynapseManual.pdf> |
| Excel1 6.38 | | | Microsoft Office | <https://www.microsoft.com/en-us/microsoft-365/excel> |
| TopScan | | | Clever Sys Inc. | <http://cleversysinc.com/CleverSysInc/csi_products/topscan-suite/> |
| ANY-maze Behavioral Tracking Software | | | Stoeling Co. | <https://www.any-maze.com/> |
| Illustrator CS5 | | | Adobe | <https://www.adobe.com> |
| R | | | R Foundation for Statistical Computing | <https://www.r-project.org/> |
| RStudio | | | RStudio | <https://www.rstudio.com/> |
| MED-PC 5 | | | Med Associates Inc | <https://www.med-associates.com/tag/med-pc-iv/> |
| MED-PC 5 Software Suite | | | This paper |  |
| Clocklab | | | Actimetrics, IL | <https://actimetrics.com/products/clocklab/> |
| pClamp/ Clampfit 11.2 | | | Molecular Devices | <http://mdc.custhelp.com/app/answers/detail/a_id/18779/~/axon%99pclamp%99-10-electrophysiology-data-acquisition-%2526-analysis-software-download> |
| Other | | | | |
| 26-gauge guide cannulas | | | P1 Technologies | C235GS-5-[spacing]/SPC GUIDE  26GA DBL (5MM PED) |
| 33-gauge injectors | | | P1 Technologies | C235IS-5/SPC INTERNAL 33GA DBL |
| Dummy cannulas | | | P1 Technologies | C235DCS-5/SPC DUMMY DBL |
| Running Disks | | | Amazon | https://www.amazon.com/Hamster-Running-Flying-Animal-Accessories/dp/B0BN3M5WN8/ref=sr_1_3?keywords=running+disk+hamster&qid=1684953974&sr=8-3 |
